# *Doublesex-*mediated regulation of insulin signaling drives sex-specific body growth

**DOI:** 10.64898/2026.06.26.734866

**Authors:** Di Guo, Si-Jie Wang, Shi-Yong Zhao, Dick R. Nässel, Cong-Fen Gao, Shun-Fan Wu

## Abstract

Most animals develop sex-biased body sizes driven by sexually divergent plasticity in nutrient-dependent growth. Prior work in *Drosophila*, implicated the sex-determination gene *transformer* (*tra*) in regulation of sex differences in body size. However, *tra* does not widely mediate sex determination across insects. Thus, the unifying molecular pathway governing female-biased sexual size dimorphism (SSD) remains poorly established in non-drosophilid insects. The rice stem borer *Chilo suppressalis*, a devastating lepidopteran crop pest, exhibits a robust SSD, offering an ideal system to dissect underlying mechanisms. We report that female-specific splice forms of the sex-determining gene *doublesex* (*Csdsx)* are master regulators of female-biased growth. Disruption of the female-specific *Csdsx* exon 3 via CRISPR knockout or RNA interference drastically reduces female body size and completely erases the SSD. *Csilp2*, encoding a key insulin-like peptide (ILP2), is selectively upregulated in late-instar female larvae, and female CsDsx proteins directly bind and activate the *Csilp2* promoter to boost transcription. Loss-of-function of *Csilp2* eliminates the SSD by suppressing female somatic overgrowth. Our results identify a novel regulatory cascade: female-specific Dsx directly stimulates insulin signaling via *Csilp2*, bridging core sex-determination circuitry and nutrient-dependent body growth control.

**Significance:** Sexual size dimorphism (SSD), widespread across insects with larger females, strongly shapes reproductive fitness. Yet reports on the molecular connection between sex determination and dimorphic growth are scarce. Using a major agricultural pest, the rice stem borer with a prominent female-biased SSD, we show that female-specific Doublesex (dsx) splice variants directly activate the promoter of Insulin-like peptide 2 in late larvae to trigger increased growth. Knocking out either gene in females abolishes the SSD. Our findings thus establish dsx as a novel link connecting sex-determination, insulin signaling and nutrient-dependent body growth.

## Introduction

Sexual size dimorphism (SSD), is a phylogenetically widespread phenomenon across the poikilothermic vertebrates and most arthropods, and arises from divergent evolutionary equilibria shaped by sex-specific pressures on natural selection (<u>Blanckenhorn, 2005</u>; <u>Stillwell et al., 2010</u>). In the majority of invertebrate species and ectothermic vertebrates, females are the larger sex, whereas male-biased SSD predominates in birds and mammals (<u>Blanckenhorn, 2005</u>; <u>Isaac, 2005</u>; <u>Monnet and Cherry, 2002</u>; <u>Teder and Tammaru, 2005</u>; <u>Webb and Freckleton, 2007</u>). This taxonomic divergence reflects distinct evolutionary optima shaped by sex-specific selection pressures. The evolutionary basis of sexual size dimorphism (SSD) centers on distinct selective mechanisms: female-biased patterns are principally driven by fecundity selection, male-biased SSD is predominantly attributed to sexual selection through male-male competition (<u>Allen et al., 2011</u>; <u>Kuntner and Elgar, 2014</u>; <u>Reeve and Fairbairn, 1999</u>).

In insects, sexual SSD arises through distinct ontogenetic mechanisms: (1) the larger sex may originate from sex-specific egg size, establishing size asymmetry at hatching (rarely examined, see e. g. (<u>Anna et al., 2013</u>)); (2) the larger sex may undergo more larval instars than the smaller sex (reviewed in (<u>Esperk et al., 2007</u>)); or (3) the larger sex exhibits more limited weight loss during metamorphosis, retaining greater post-metamorphic mass (<u>Molleman et al., 2011</u>; <u>Testa et al., 2013</u>; <u>Vendl et al., 2018</u>). Most research has focused on resolving whether the larger size in one sex is primarily attained through longer developmental periods or faster growth of the juveniles. In insects, accumulating evidence demonstrates that larvae of the larger sex tend to grow for a longer time than those of the smaller sex (<u>Stillwell and Davidowitz, 2010</u>; <u>Tammaru et al., 2010</u>; <u>Teder, 2014</u>), although sexually dimorphic growth rates have also been documented (<u>Blanckenhorn et al., 2007</u>; <u>Knapp, 2014</u>; <u>Rohner et al., 2016</u>; <u>Vendl et al., 2018</u>).

In *Drosophila*, the sex-determination gene *Sex-lethal* (*Sxl*) regulates SSD by controlling body growth during larval development through its action in female-specific neurons of the nervous system (<u>Sawala and Gould, 2017</u>). During the larval stage, *Sxl* selectively promotes the growth rate of larval tissues, but not imaginal disc tissues that give rise to adult tissues (<u>Sawala and Gould, 2017</u>). Another gene in the sex-determination pathway, *transformer* (*tra*), contributes to SSD regulation in *Drosophila* (<u>Rideout et al., 2016</u>). *tra* is downstream of *Sxl* in the sex-determination cascade, and regulates cell size through cell-autonomous effects, but also indirectly via action in the female fat body to stimulate neurosecretory cells in the brain to secrete insulin-like peptides (ILPs), thereby promoting general growth and increased body size (<u>Rideout et al., 2016</u>).

In other insects, further sex-determination genes, such as *doublesex* (*dsx*) have been implicated in sex-specific growth. In the beetle *Onthophagus taurus*, the tissue-specific expression of sex-specific *dsx* isoforms promotes or suppresses the condition-dependent growth of horns (<u>Kijimoto et al., 2012</u>). Intriguingly, *dsx* not only regulates sexual dimorphism in horn development but also modulates the nutrient sensitivity of horn growth. In the stag beetle, *Cyclommatus metallifer*, the extraordinary development of male mandibles depends on the male-specific splice form of *dsx* conferring sex-specific responsiveness to juvenile hormone (JH) signaling. Conversely, in females, differential splicing of *dsx* prevents JH from activating the mandibular growth pathway (<u>Gotoh et al., 2014</u>).

The rice stem borer, *Chilo suppressalis* Walker (Lepidoptera: Crambidae), ranks among the most devastating pests in global rice production, with its larval stage inflicting significant economic losses on rice crops (<u>Dutta and Roy, 2022</u>). Adults exhibit high fecundity, and a single female lays 100–300 eggs. Newly hatched larvae feed gregariously within leaf sheaths, while second instar bore into rice stems. This feeding behavior induces damaged tillers (<u>Dutta and Roy, 2022</u>). Notably, conspicuous sexual size dimorphism is observed in both adults and pupae of *C. suppressalis*. However, the genetic and physiological mechanisms that account for the larger female body size remain unclear. Here, using *C. suppressalis* as a model insect, we establish for the first time that *dsx* functions as a key regulator of SSD. Furthermore, we demonstrate that *dsx* influences sex-specific somatic growth in females through a novel regulatory axis involving direct modulation of the insulin-signaling pathway. We show that Dsx binds to and activates the promoter of one of the insulin genes. This gene encodes insulin-like peptide 2, a peptide that increases its expression in females during the sex-specific growth and whose knockdown eliminates the SSD.

## Results

### *C. suppressalis* displays a sexual size dimorphism at the late larval, pupal, and adult stages

The *C. suppressalis* embryo hatches into a larva that commonly develops through 6 instars (L1 to L6) (Table S1). However, some individuals develop into 7th or 8th instars, and even 9th instars, but most of these die without pupating (<u>Luo et al., 2016</u>). The insect then enters a long non-feeding pupal phase, which ends with eclosion of the adult moth. Final body size in *C. suppressalis,* as in in other insects, is determined by the availability of food and feeding rate, the larval growth rate, and the duration of the growth period (<u>Shingleton and Vea, 2023</u>). However, no significant difference in egg size is observed between sexes (Fig. S1A-B). To understand when sexual size dimorphism (SSD) first appears during development, we compared the growth curves of male and female larvae. Individual examination of larvae at precise times after hatching revealed that the sizes are similar in L1 to L4 males and females and that a SSD first becomes detectable during L5 and then increases until L6 larvae (Fig. 1A-C). The female pupae and adult moths are approximately 30% larger than their male counterparts (Fig. 1D-F). Morphometric comparisons between adult sexes revealed that females exhibit approximately 40% greater body mass than males. Furthermore, measurements of wing length, body length, as well as abdominal and thoracic width demonstrated significantly larger dimensions across all morphological parameters in females compared to males (Fig. 1G-N). Taken together, these results indicate that SSD is established at L5 larval stage of the rice stem borer.

**Fig. 1.**
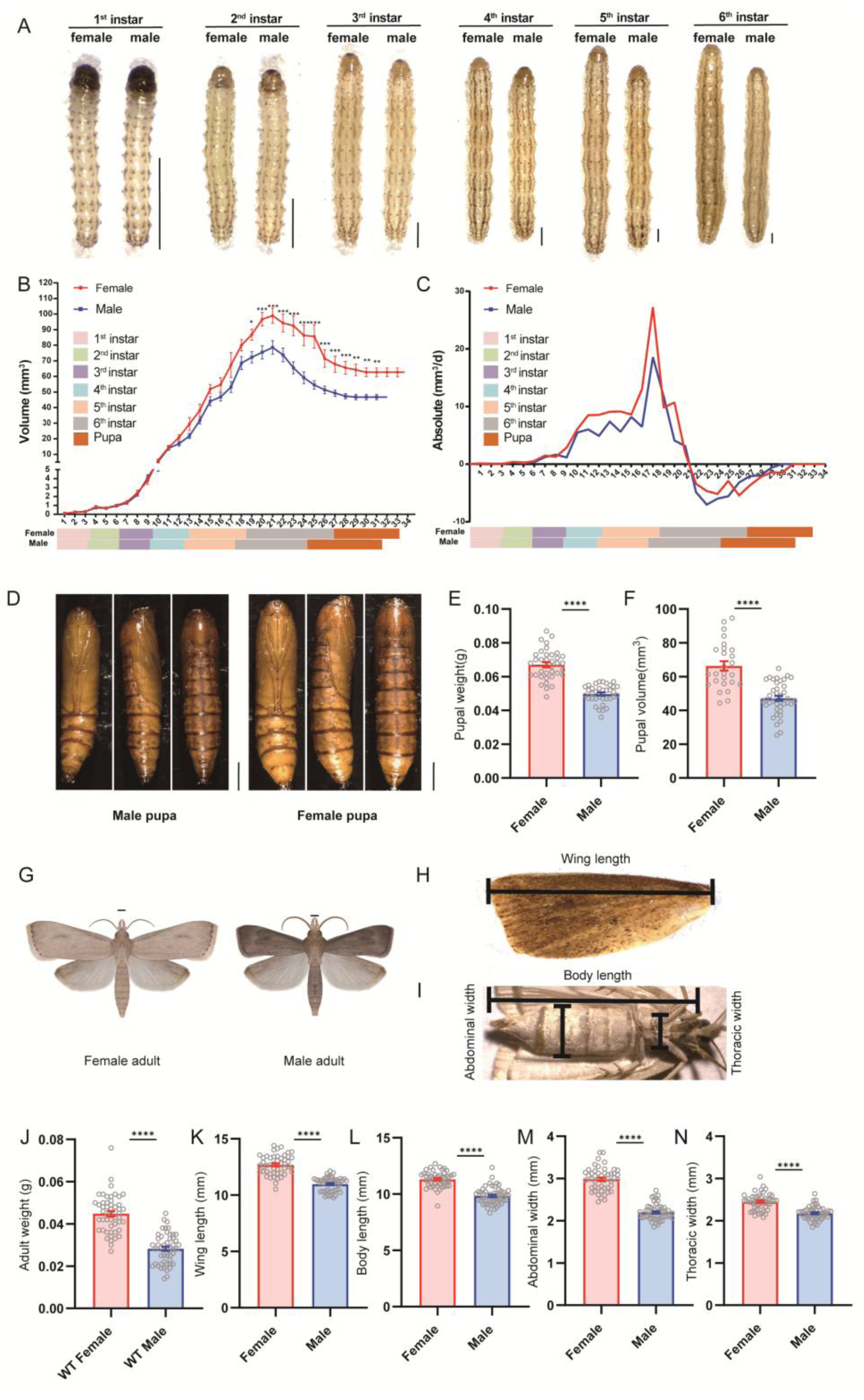
Sexual size dimorphism in *C. suppressalis*. (A) Morphological features of female and male larvae across instar stages in *C*. *suppressalis*. Shown are L1 to L6 larval stages. Scale bar = 1 mm. (B) Growth curves of male and female larvae throughout larval development, showing raw volumetric measurements at each time point. Data were analyzed using one-way ANOVA with Tukey’s multiple comparisons test, (*, *p* <0.05; ***, *p* < 0.001; ****, *p* < 0.0001; ns, not significant). (C) Absolute growth rates across larval developmental stages. (D) Dorsal, lateral, and ventral views of female and male pupae. (E) Comparison of pupal weight differences between females and males. (F) Comparison of pupal volume differences between females and males. Data are expressed as mean ± SEM, Student’s t-test, (*, *p* <0.05; ***, *p* < 0.001; ****, *p* < 0.0001; ns, not significant). Scale bar = 1 mm. (G) Morphological features of female and male adults. (H-I) Schematic diagram of morphological measurements for female and male adults. (J-N) Comparison of body weight, wing length, body length, abdominal width, and thoracic width between female and male adults. Data are expressed as mean ± SEM; Student’s t-test, (*, *p* <0.05; ***, *p* < 0.001; ****, *p* < 0.0001; ns, not significant).

**Table S1.** Median duration of development (days) for different instar stages.

|  | Instar number |  |  |  |  |  |  |
| --- | --- | --- | --- | --- | --- | --- | --- |
|  | 1<sup>st</sup> | 2<sup>nd</sup> | 3<sup>rd</sup> | 4<sup>th</sup> | 5<sup>th</sup> | 6<sup>th</sup> | 7<sup>th</sup> |
| Female | 3 (2-5) | 3 (2-8) | 3 (2-6) | 3 (2-5) | 5 (4-11) | 9 (4-12) | 10 (8-11) |
| male | 3 (2-3) | 3 (2-4) | 3 (3-3) | 3 (3-5) | 5 (3-8) | 7 (4-10) | 7 (6-8) |

Numbers in brackets represent ranges.

**Fig. S1.**
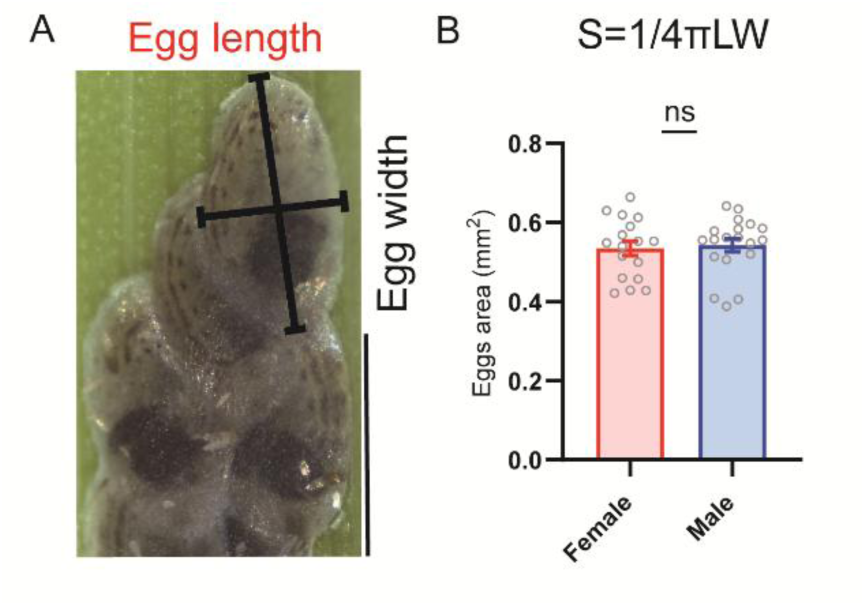
Comparison of egg size between female and male *Chilo suppressalis*. (A) Schematic diagram of egg length and width (black lines) measurements in *Chilo suppressalis*. (B) Comparison of egg area between female and male eggs (S=surface area of the egg). For sex determination, eggs were collected immediately prior to hatching and DNA was extracted for molecular analysis (see methods).

**Fig. S2.**
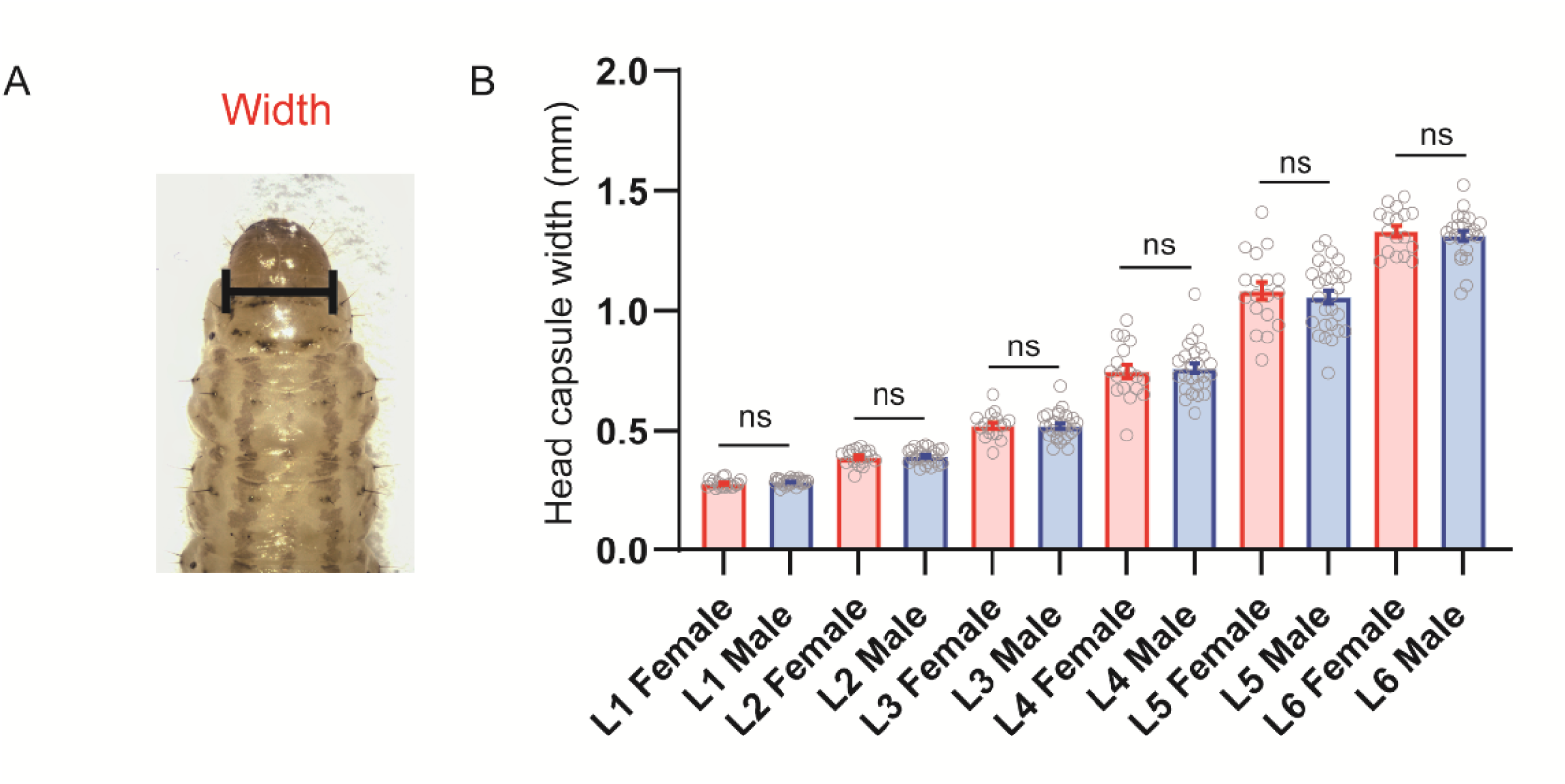
Head capsule width of female and male larvae at different instar stages in *Chilo suppressalis*. (A) Image of head capsule width measurements in *Chilo suppressalis*. (B) Head capsule width of female and male larvae at different instar stages. Data are expressed as mean ± SEM, student’s t-test, (*, *p* <0.05; ***, *p* < 0.001; ****, *p* < 0.0001; ns, no significant).

### *Doublesex* regulates sex differences in body size of the rice stem borer

In *C. suppressalis,* the *Csdsx* gene gives rise to three splice forms, one male-(*Csdsx^M^*) and two female-specific forms (*Csdsx^F1^ and Csdsx^F2^*) (Fig. 2A). We analyzed the expression of *Csdsx* splice forms at different developmental stages of both sexes (No difference in body size between females and males is observed prior to the third-instar larval stage) (Fig. S3A) and in different body parts of male and female larval instars (Fig. S3B). These data show that relative expression levels of *Csdsx* during L1 to L4 are not significantly different in males and females. Interestingly, however, thereafter the expression of the exons *Csdsx^E3^*and *Csdsx^E4^* (representing two female-specific forms *Csdsx^F1^ and Csdsx^F2^*) is higher in the female larvae than that of the exon *Csdsx^E5^* (representing male-specific forms *Csdsx^M^*) in the male larvae (Fig.S3A). As a result, *Csdsx^E3^* and *Csdsx^E4^* displayed significantly higher expression in the head, epidermis, Malpighian tubules and fat body of females than *Csdsx^E5^* in corresponding male tissues (Fig. S3B).

**Fig. 2.**
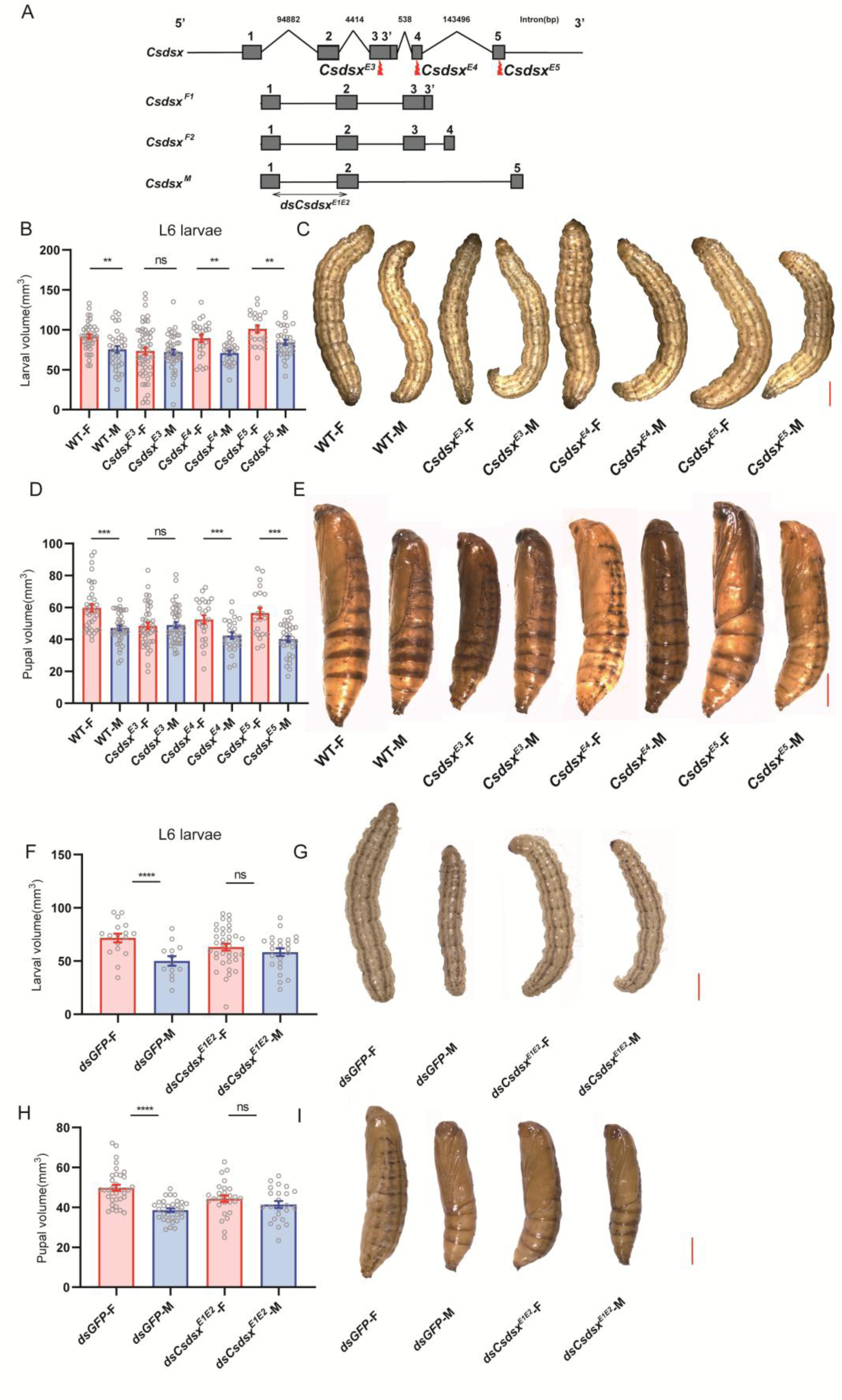
The sex determination gene *doublesex* (*Csdsx*) regulates sex differences in body size. (A) Characterization of *C. suppressalis dsx* gene (*Csdsx*). The three different sex-specific transcripts in males (*Csdsx^M^*) and females (*Csdsx^F1^* and *Csdsx^F2^*) are shown. Exons are indicated by boxes and introns by lines. (B) Quantitative analysis of larval volume <u>[(calculated according to the *Drosophila* pupal-volume formula: V=1/6π(L)(W)^2^</u> (<u>Rideout et al., 2016</u>)]. In wild-type and mutant male and female 6^th^ instar (L6) larvae. (C) Images of wild-type and mutant male and female larvae (L6). (D) Quantitative analysis of pupal volume in wild-type and mutant male and female pupae. The female pupal volume is approximately 30% larger than males in wild-type strains. (E) Images of wild-type and mutant male and female pupae. (F) Quantitative analysis of female and male larval (L6) volume after *dsRNA* (*Csdsx*) treatment in late L5. The dsRNA targeted the different exons E3-E5 shown in Fig. 2A. (G) Images of female and male larval volume after *dsRNA*. (H) Quantitative analysis of female and male pupal volume after feeding of dsRNA in L6. (I) Images of female and male pupal volume post *dsRNA*. Student’s t-test, (*, *p* <0.05; ***, *p* < 0.001; ****, *p* < 0.0001; ns, no significant).

We therefore asked whether the size might be regulated by *Csdsx* and measured volume and weight in *Csdsx* mutant animals (Fig 2A). We found that body size was significantly reduced in *Csdsx^E3^* mutant females compared to the wild-types (Fig 2D and E). Thus, while wild-type females are larger than males, *Csdsx^E3^* mutant females are not. This suggests that *Csdsx^E3^* contributes to establishing SSD in *C. suppressalis.* Importantly body size was unchanged in *Csdsx* mutant males (Fig 2D and 2E). We also monitored the size of L6 larvae in *Csdsx* mutant animals, and found a similar abolishment of SSD in *Csdsx^E3^*animals, but not in mutants of the other splice exons (Fig 2B and 2C). To further validate whether female-specific body size is regulated by *Csdsx*, we performed targeted gene silencing during the critical developmental stage of the SSD in *C. suppressalis*. Early fifth-instar larvae were subjected to *dsx* knockdown (RNAi) via dsRNA feeding (Fig 2A and S4A-B). We observed that an abolishment of the SSD occurred in knockdown animals during the peak SSD divergence phase seen in wild-type individuals at the initial sixth-instar stage (Fig 2F and G). Importantly, we found that *dsx*-silenced pupae exhibited abolished sexual dimorphism in pupal volume compared to controls (Fig 2H and I). Taken together, these results suggest that male-female differences in body size originates in part by the action of *dsx* (*Csdsx^E3^*) in females starting at L6, thus establishing a new role for *dsx* in the regulation of sex differences in body growth.

**Fig. S3.**
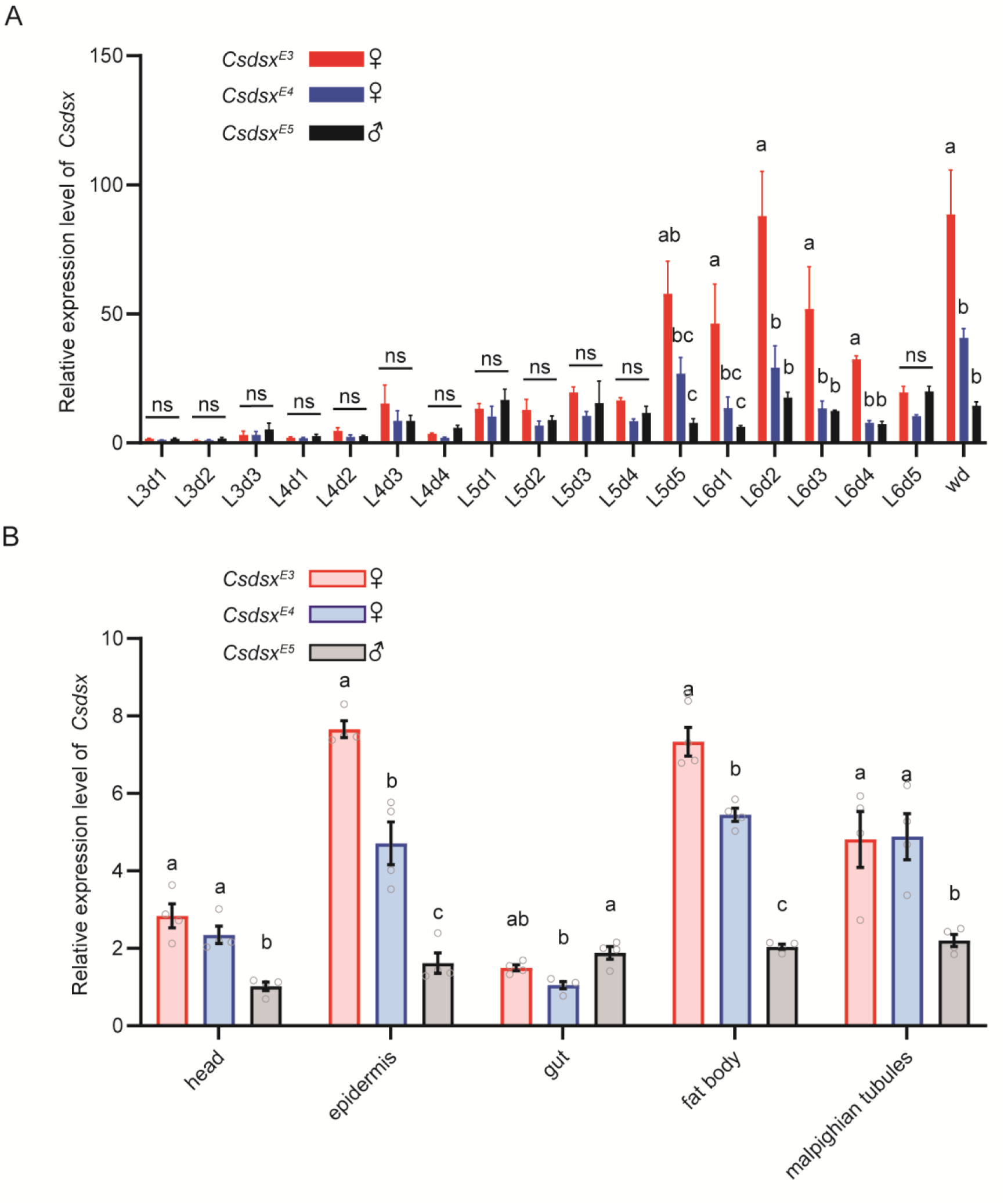
Characterization of the *Chilo suppressalis dsx* transcript. (A) Expression analysis of *Csdsx* in different developmental stages of male and female larvae. Expression levels of *Csdsx* were measured based on exons 3 and 4 (E3 and E4; female-specific splice variants) and exon 5 (E5, male-specific splice variant). Developmental stage annotations: L3d1: Day 1 of the 3rd instar larval stage (and so on); wd: Wandering stage (prepupal phase). (B) Tissue-specific expression analysis of *Csdsx* in larvae. In female larval tissues, the relative expression levels of exons 3-4 were measured; in male larval tissues, the relative expression level of exon 5 was measured. Statistical Annotation: Different lowercase letters (a, b, c…) indicate significant differences between groups determined by one-way ANOVA with Tukey’s multiple comparisons test (*p* < 0.05).

**Fig S4.**
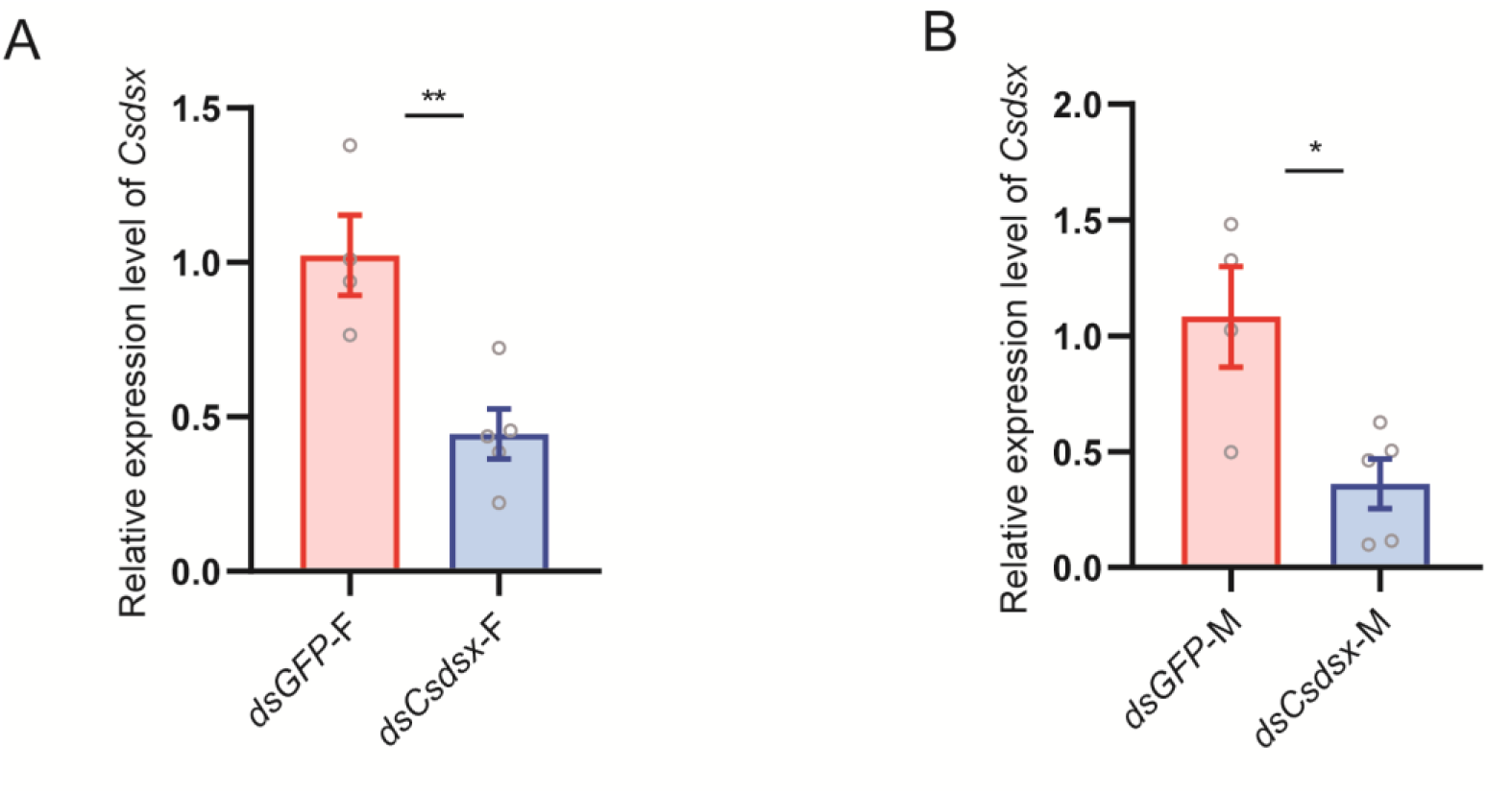
Silencing efficiency of *doublesex* gene after dsRNA feeding. (A-B) Validation of *dsx* silencing efficiency by qRT-PCR. Student’s t-test, (*, *p* <0.05; ***, *p* < 0.001; ****, *p* < 0.0001.

### Transcriptomic screening identifies *Csilp2* as the sex-specific insulin effector governing female-biased body growth

To identify potential genes mediating the sexual size dimorphism in *C. suppressalis* larvae, differential gene expression analysis was performed on transcriptome sequencing data. Using filtering criteria (q-value < 0.05; |Log₂FC| > 1), we compared gene expression profiles between sexes across three developmental stages: fourth-instar, fifth-instar, and sixth-instar larvae. Cross-stage comparative analysis revealed 213 overlapping differentially expressed genes (DEGs) between fourth- and fifth-instar larvae; 190 overlapping DEGs between fourth- and sixth-instar larvae; and 546 overlapping DEGs between fifth- and sixth-instar larvae. Notably, 182 core DEGs were conserved across all three developmental stages (Fig. 3A). Building on our findings that pronounced sexual size dimorphism emerges in *C. suppressalis* during the fifth-instar larval stage, Gene Ontology enrichment analysis of female-biased differentially expressed genes revealed significant functional enrichment in three core biological domains: signal transduction pathways, metabolic regulation, and cellular motility coupled with structural maintenance (Fig. 3B). Therefore, we focused on sexually dimorphic genes in fifth-instar *C. suppressalis* larvae, revealing significant upregulation of insulin signaling pathway components in females.

**Fig. 3.**
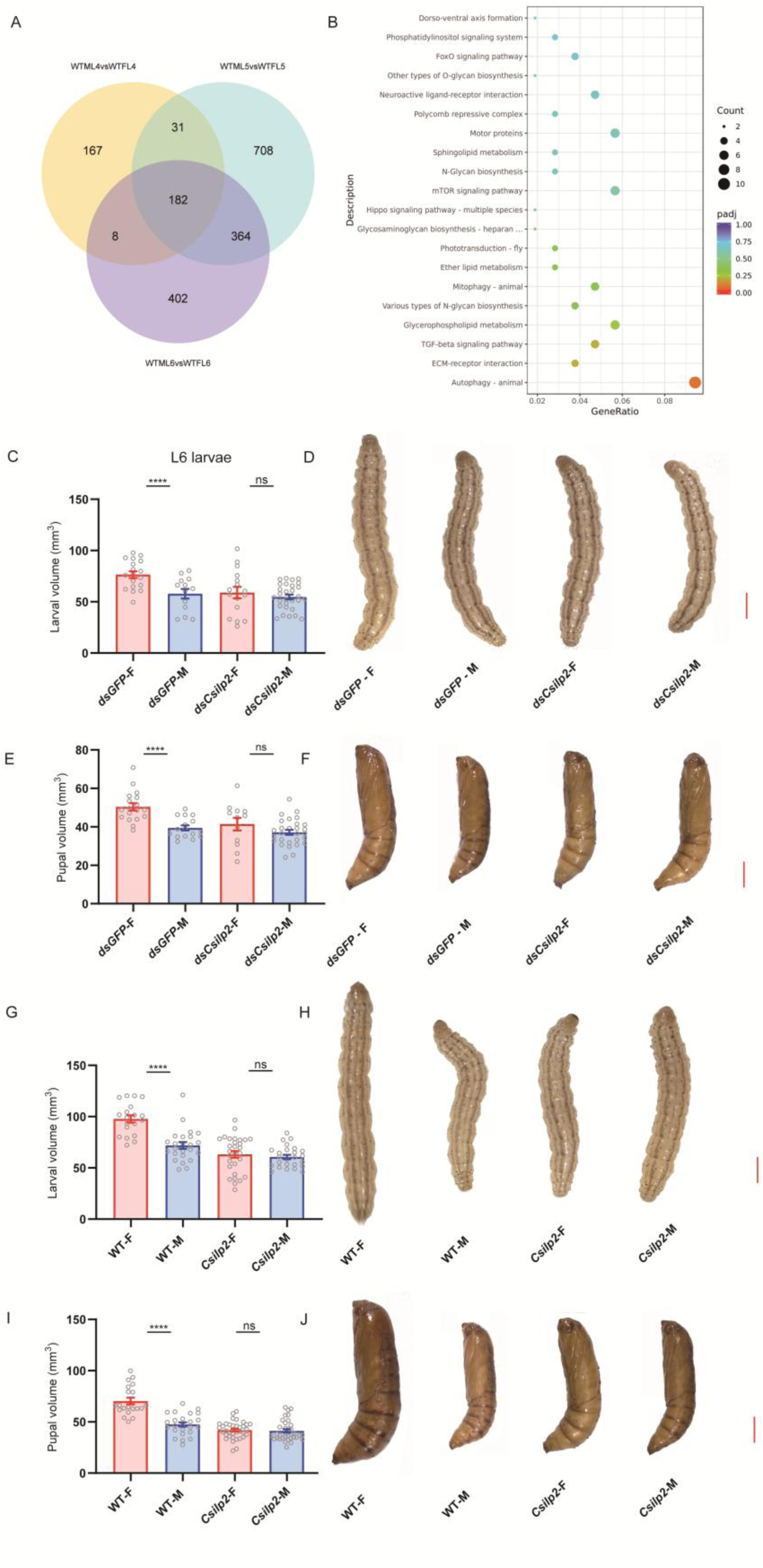
The insulin-like peptide 2 gene, *Csilp2,* regulates sex differences in body size. (A) Venn diagram analysis of the numbers of differential gene expression between male and female larvae of *C. suppressalis* across developmental stages. WTML4: Wild-type male 4th instar larvae; WTFL4: Wild-type female 4th instar larvae (and so on) (B) Y-axis indicates KEGG pathways, and X-axis represents gene ratio. KEGG pathway enrichment is considered significant at padj (p-value adjusted by the Benjamini-Hochberg method (FDR)) < 0.05. (C-D) Quantitative analysis and images of female and male larval volume after feeding with dsRNA at early 5th instar larva stage. (E-F) Quantitative analysis of female and male pupal volume after feeding with *Csilp2 Csilp2* dsRNA at early 5th instar larva stage. (G-H) Quantitative analysis of body size and images of female and male larvae in wild-type and *Csilp2* mutant strains. (I-J) Quantitative analysis of body size of female and male pupae in wild-type and *Csilp2* mutant strains. Statistical annotation: Data are presented as mean ± SEM. Student’s t-test. (*, *p* <0.05; ***, *p* < 0.001; ****, *p* < 0.0001; ns, not significant). Scale bar: 2 mm.

From the *C. suppressalis* transcriptome, we identified and cloned six genes encoding insulin-like peptides (ILPs), *Csilp1 - 6*. Multiple sequence alignment revealed highly conserved amino acid sequences in the A- and B-chains of these ILPs, whereas the C-peptide and signal peptide domains exhibited lower conservation. Phylogenetic analysis demonstrated evolutionary conservation between *C. suppressalis* ILPs and those of the lepidopteran model, the silk moth *Bombyx mori* (Fig. S5A and S8). Interestingly, we found no IGF-like ILP in this species in contrast to ILP6 in *Drosophila* and the IGF-like peptide in *Bombyx* (Fig. S5A and B). Next, we asked whether expression of any of these genes was differentially altered during development in females compared to males. Notably, *Csilp2* expression in female larvae exceeded that in males during the fifth instar (Fig. S5B, S6A and S7), whereas no significant differences between sexes were observed for other *Csilp transcripts (Csilp1 and Csilp3 – 6,* or insulin signaling pathway genes such as the ILP receptor (InR) and FOXO (Fig. S7). This indicates that *Csilp2* likely plays a regulatory role in late-stage larval growth in *C. suppressalis*.

To investigate the role of ILP signaling genes in *C. suppressalis* development, we silenced *Csilp2* and its receptor *Csinr* (by feeding dsRNA) at early 5th instar larva stage (Fig. S9A and B, S11A and B). *Csilp2* and *Csinr* knockdown significantly reduced larval growth rates (Fig. S10A). Not surprisingly, *Csinr* silencing induced approximately 75% larval mortality, whereas *Csilp2* knockdown had no impact on survival (Fig. S10B). Thus, we focused on analyzing body size effects following *Csilp2* silencing. Both pupal volume measurements and sixth-instar larval assessments revealed diminished SSD in *Csilp2*-silenced individuals (Fig. 3C, D, E and F). To further investigate the role of *Csilp2* in *C. suppressalis* development, we generated *Csilp2* knockout lines using CRISPR/Cas9 (Fig. S9D and E). These *Csilp2* mutants exhibited significant body size reduction in both larvae and pupae of both sexes, accompanied by abolition of the SSD (Fig. 3G, H, I and J). Additionally, *Csilp2* knockout substantially prolonged the duration of larval developmental (Fig. S9C). These results demonstrate that *Csilp2* is essential for normal growth in *C. suppressalis* of both sexes, with the mutation exerting a more pronounced impact on female body size and consequently eliminating SSD. We also generated *Csinr* mutant lines (Fig. S11C and D). The *Csinr* mutant larvae exhibited severe growth retardation, failed to complete the larval developmental cycle, and ultimately were unable to pupate (Fig. S10C and D). Taken together our results demonstrate that the *Csilp2* branch of the insulin-like signaling pathway not only plays a critical role in overall growth regulation, but also exerts sex-specific effects on the SSD.

**Fig. S5.**
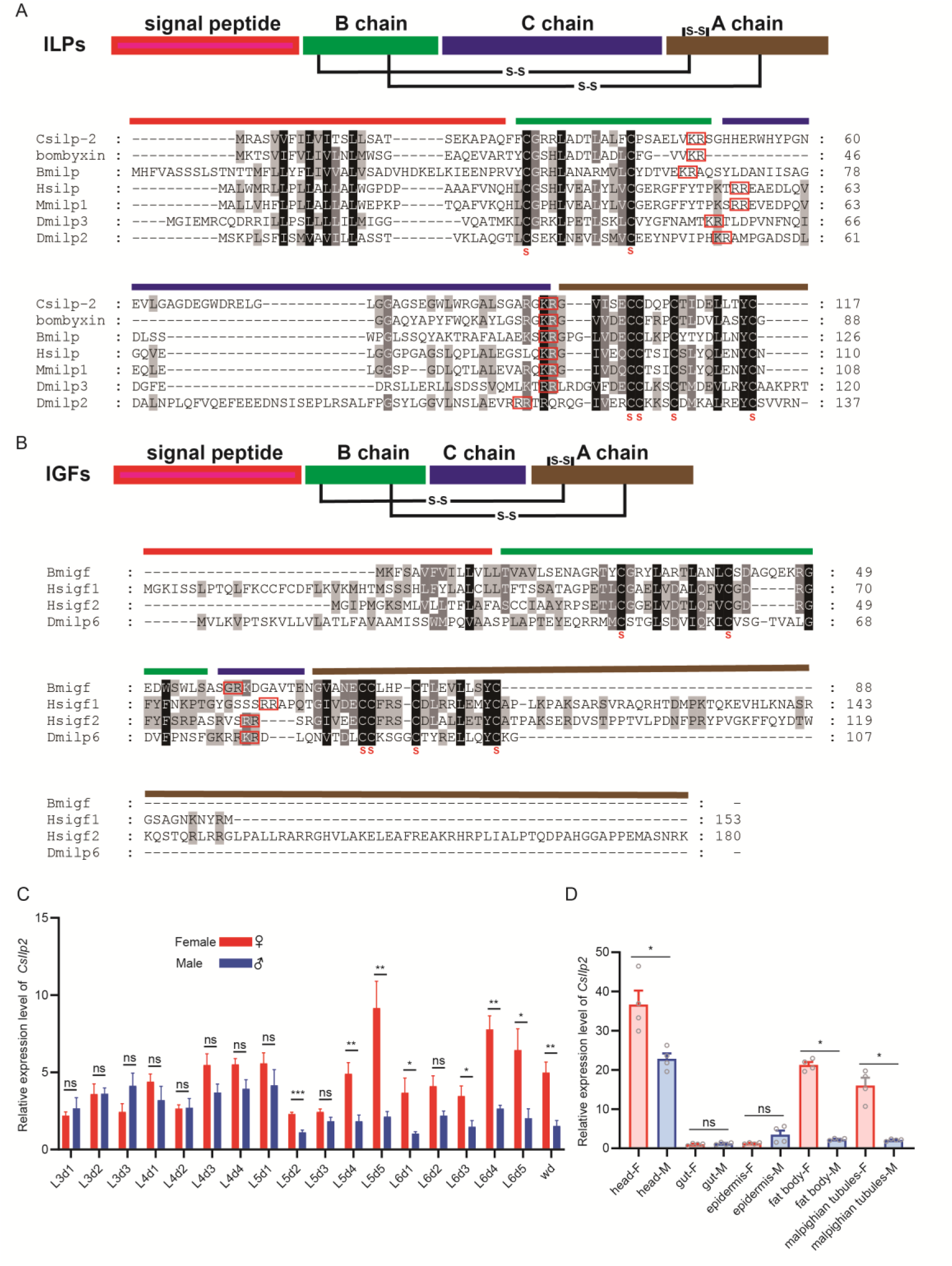
Amino acid sequence alignment of ILPs and Insulin-like Growth Factors and expression analysis of *Csilp2*. (A and B) Multiple sequence alignment of ILPs and IGFs proteins. Note that no IGF-like peptide was detected in *C. suppressalis*. Species included: Human (*Homo sapiens*, Hs), mouse (*Mus musculus*, Mm), fruit fly (*Drosophila melanogaster*, Dm) and silkworm (*Bombyx mori*, Bm). Sequence annotations: Amino acid positions are labeled on the right side of the sequences. Fully conserved residues in orthologous sequences are displayed as white text on black background. Conservatively substituted residues are highlighted with gray shading. Functional domain division: The sequences are divided into four functional regions: signal peptide (red), B-chain (green), C-peptide (blue), and A-chain (brown), arranged sequentially from the N-terminus. Disulfide bonds (S-S) are marked between the corresponding cysteine residues. (C) *Csilp2* expression across developmental stages. Developmental stage annotations: L3d1: 3rd instar day 1; wd: Wandering stage (prepupal phase). (D) Tissue-specific expression profiling of *Csilp2* in larvae. Tissue samples analyzed: Head, gut, fat body, Malpighian tubules, epidermis. Statistical annotation: Data are presented as mean ± SEM. Student’s t-test. (*, *p* <0.05; ***, *p* < 0.001; ****, *p* < 0.0001; ns, no significant).

**Fig S6.**
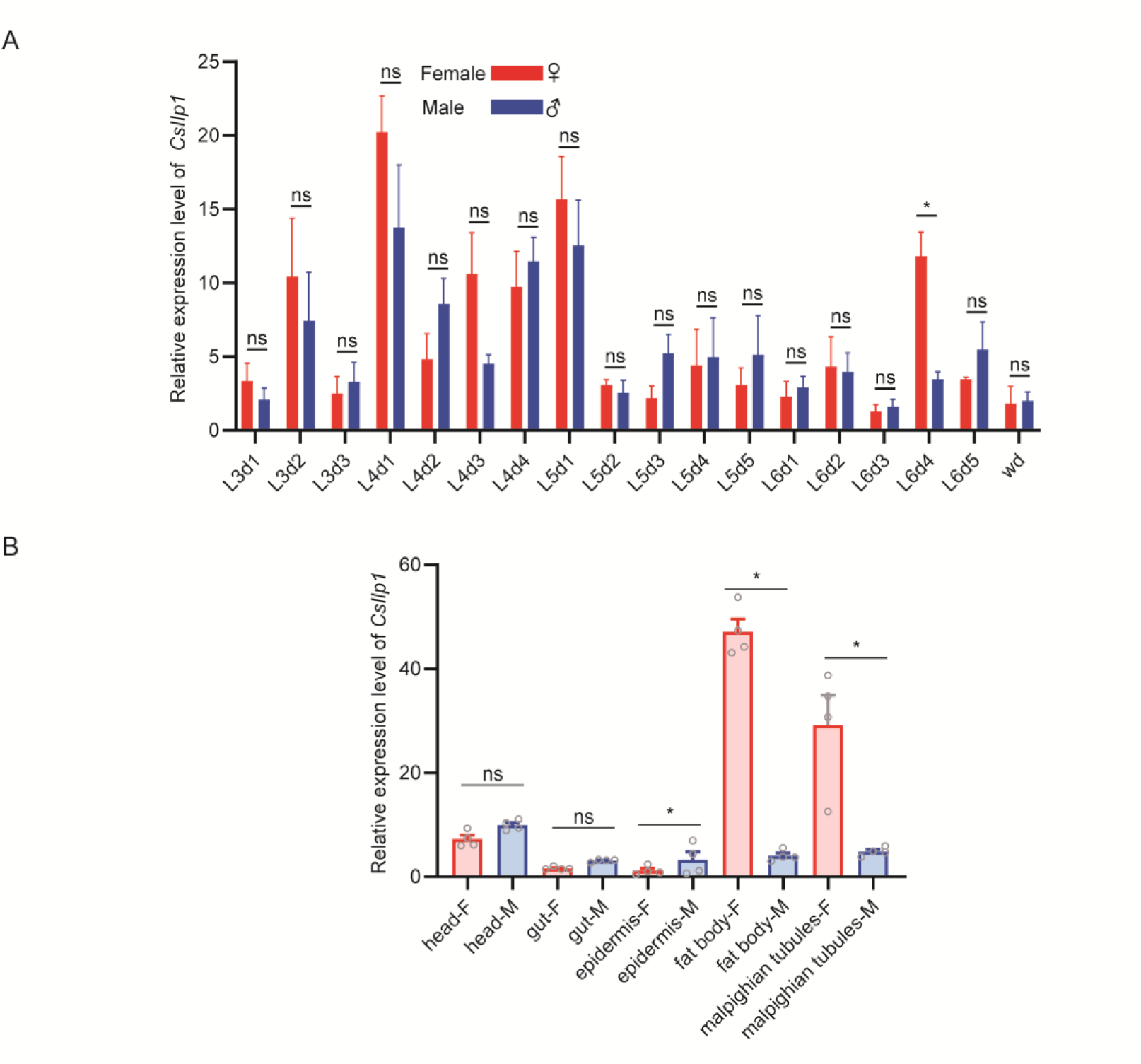
Expression analysis of *Csilp1* in different developmental stages and tissues. (A) *Csilp1* expression across developmental stages shows only a minor difference between the sexes in stage L6d5. Developmental stage annotations: L3d1: 3rd instar day 1; wd: Wandering stage (prepupal phase). (B) Tissue-specific expression profiling of *Csilp1* in larvae displays higher levels in female fat body and Malpighian tubules. Tissue samples analyzed: Head, gut, fat body, Malpighian tubules, epidermis. Statistical annotation: Data are presented as mean ± SEM. Student’s t-test. (*, *p* <0.05; ***, *p* < 0.001; ****, *p* < 0.0001; ns, no significant).

**Fig. S7.**
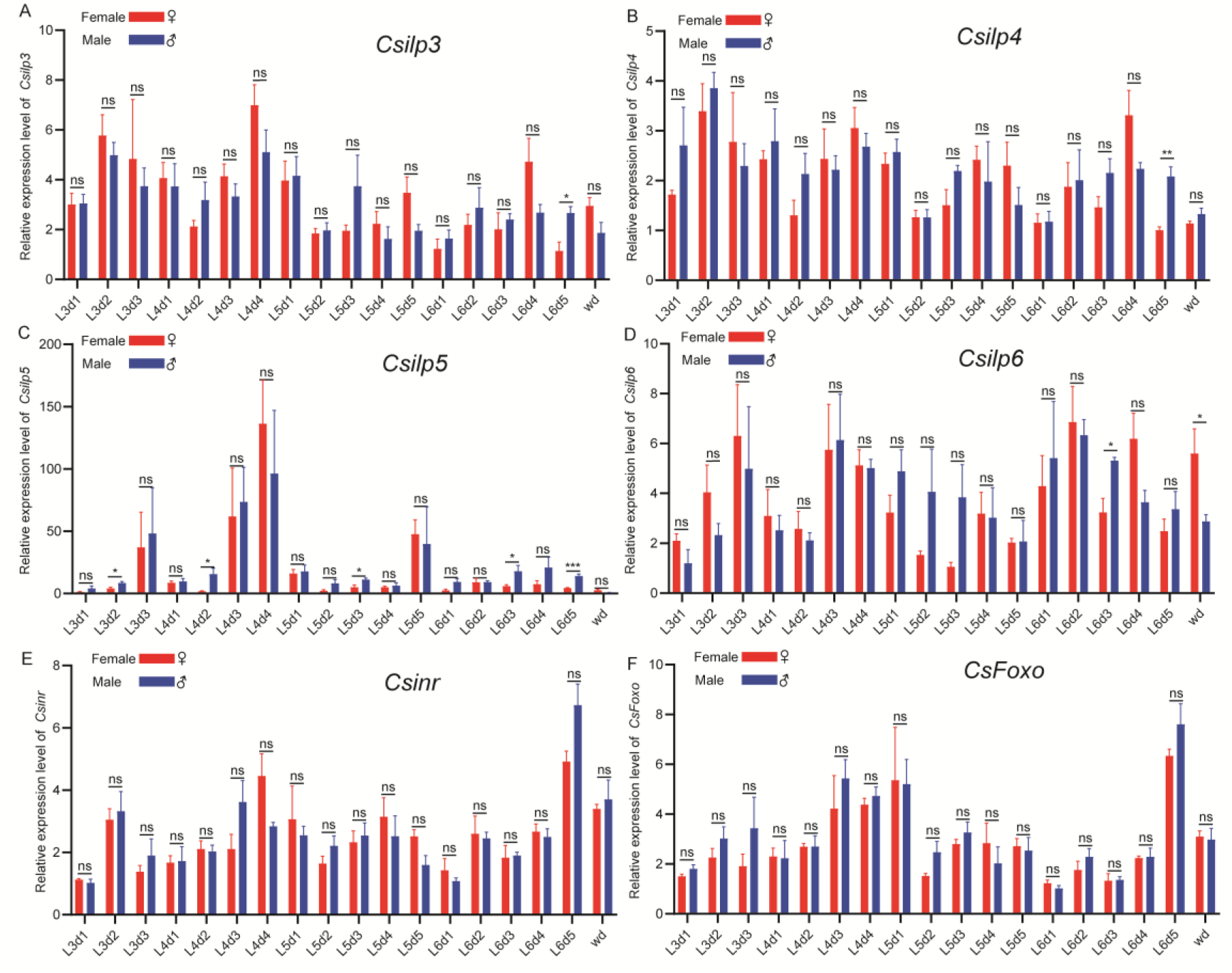
Expression levels of insulin signaling pathway genes in larvae of *Chilo suppressalis* display no sexual dimorphism during development. (A-D) *Csilp3 - 6* expression across different developmental stages. Developmental stage annotations: L3d1: 3rd instar day 1; wd: Wandering stage (prepupal phase). (E-F) *Csinr* (insulin receptor) and *CsFoxo* expression analysis in male and female larvae. Statistical annotation: Data are presented as mean ± SEM. Student’s t-test. (*, *p* <0.05; ***, *p* < 0.001; ****, *p* < 0.0001; ns, no significant).

**Fig S8.**
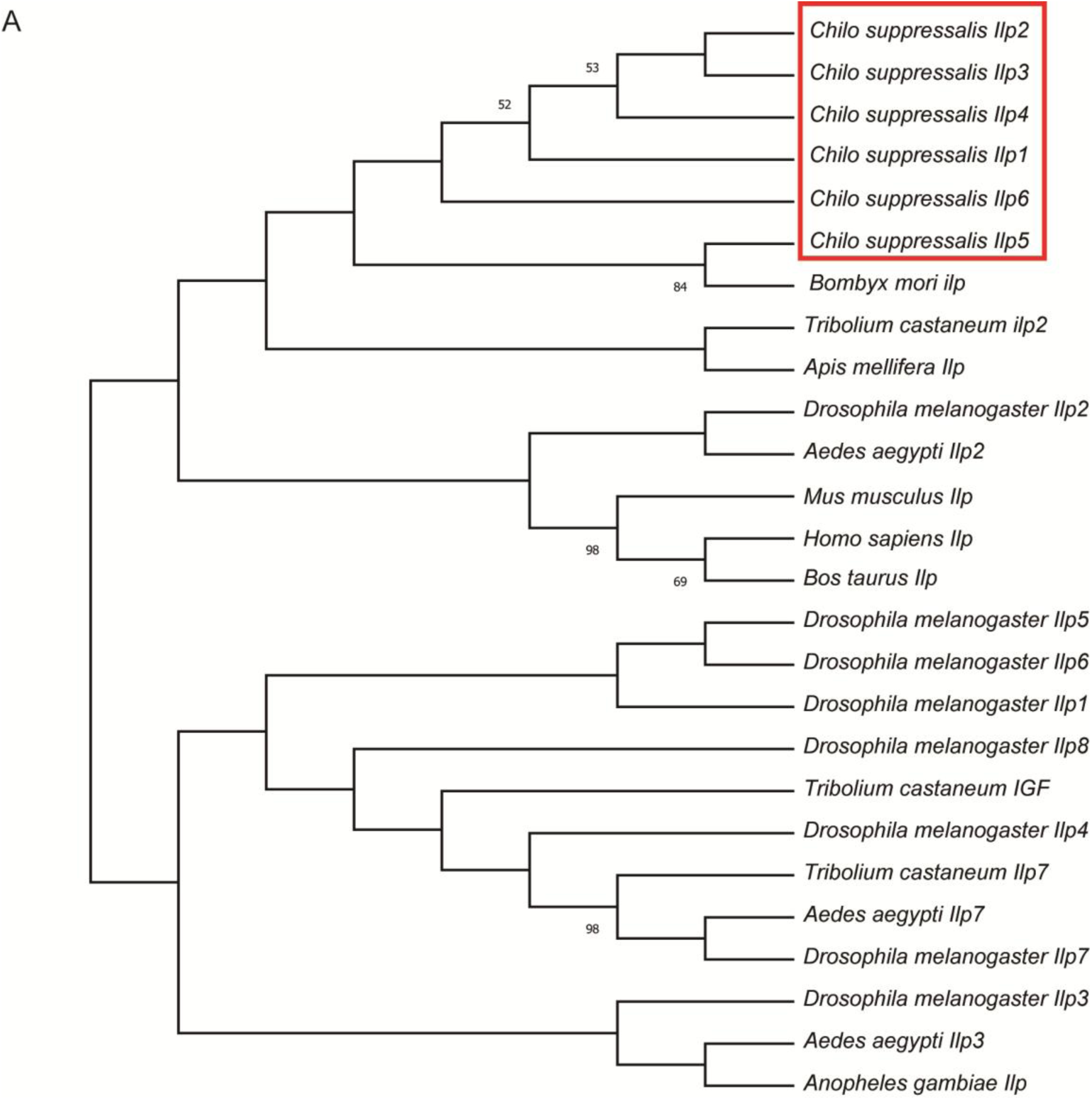
Phylogenetic tree of ILPs proteins. (A) The phylogenetic tree was constructed using the Maximum Likelihood Method in MEGAX based on multiple sequence alignment of ILP amino acid sequences from insects and mammalian species. Topological confidence was evaluated with 1,000 bootstrap replicates, and branch node values indicate bootstrap support percentages (range: 0–100%). Accession numbers for the protein sequences: *Anopheles gambiae*: XP_314565.3; *Tribolium castaneum*: XP_001814181.1, XP_064212535.1, XP_015840916.1; *Bombyx mori*: XP_012548888.1; *Apis mellifera*: NP_001171374.1; *Drosophila melanogaster*: NP_648359.1, NP_524012.1, NP_648360.2, NP_648361.1, NP_996037.2, NP_570000.1, NP_570070.1, NP_648949.2; *Mus musculus*: NP_032412.3; *Homo sapiens*: NP_001278826.1; *Bos taurus*: NP_001172055.1; *Aedes aegypti*: XP_001662922.1, ABI64117.2, XP_001657487.1.

**Fig. S9.**
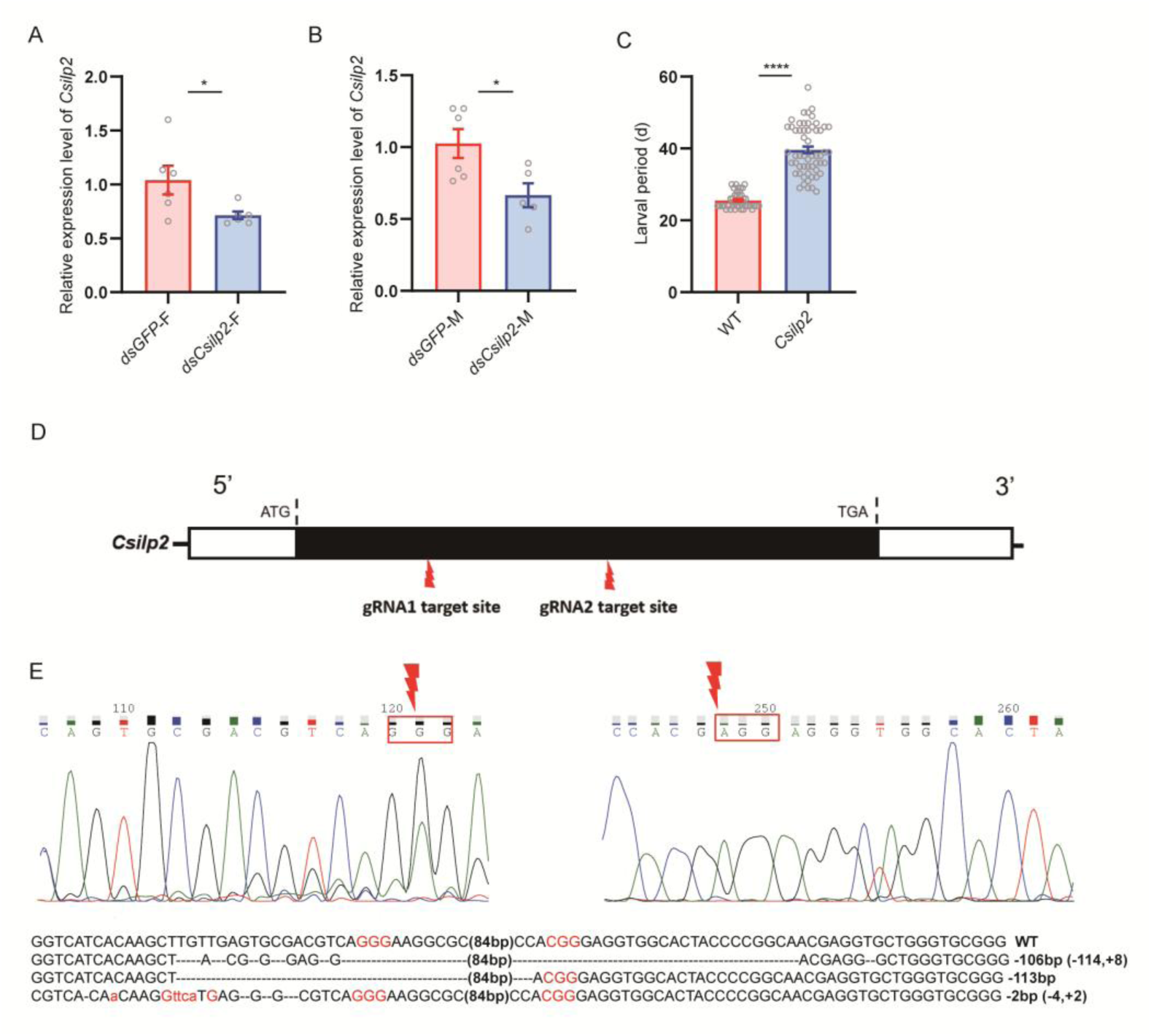
Validation of RNA interference and mutants of *Csilp2*. (A and B) Validation of *Csilp2* RNAi silencing efficiency by q-PCR in L5 larvae. (C) Duration of larval period of *Csilp2* mutants. (D) Schematic diagram of gene sequence and two sgRNA target sites. Representative sequence chromatogram of direct sequencing of PCR products of G0 individuals indicating that the two sgRNA sites were successfully cleaved. Green boxes indicate the protospacer adjacent motif (PAM) sequence. The position of the red lightning indicates the Cas9 protein cleavage site. (E) PAM sequences are highlighted in red. *Csilp2* mutant sequences were confirmed by cloning and sequencing. Dashed lines represent the deleted bases, and inserted bases are displayed in lower case. The net change in length is marked at the right of each sequence (−, deletion; +, insertion). Statistical annotation: Data are presented as mean ± SEM. Student’s t-test. (*, *p* <0.05; ***, *p* < 0.001; ****, *p* < 0.0001; ns, no significant).

**Fig. S10.**
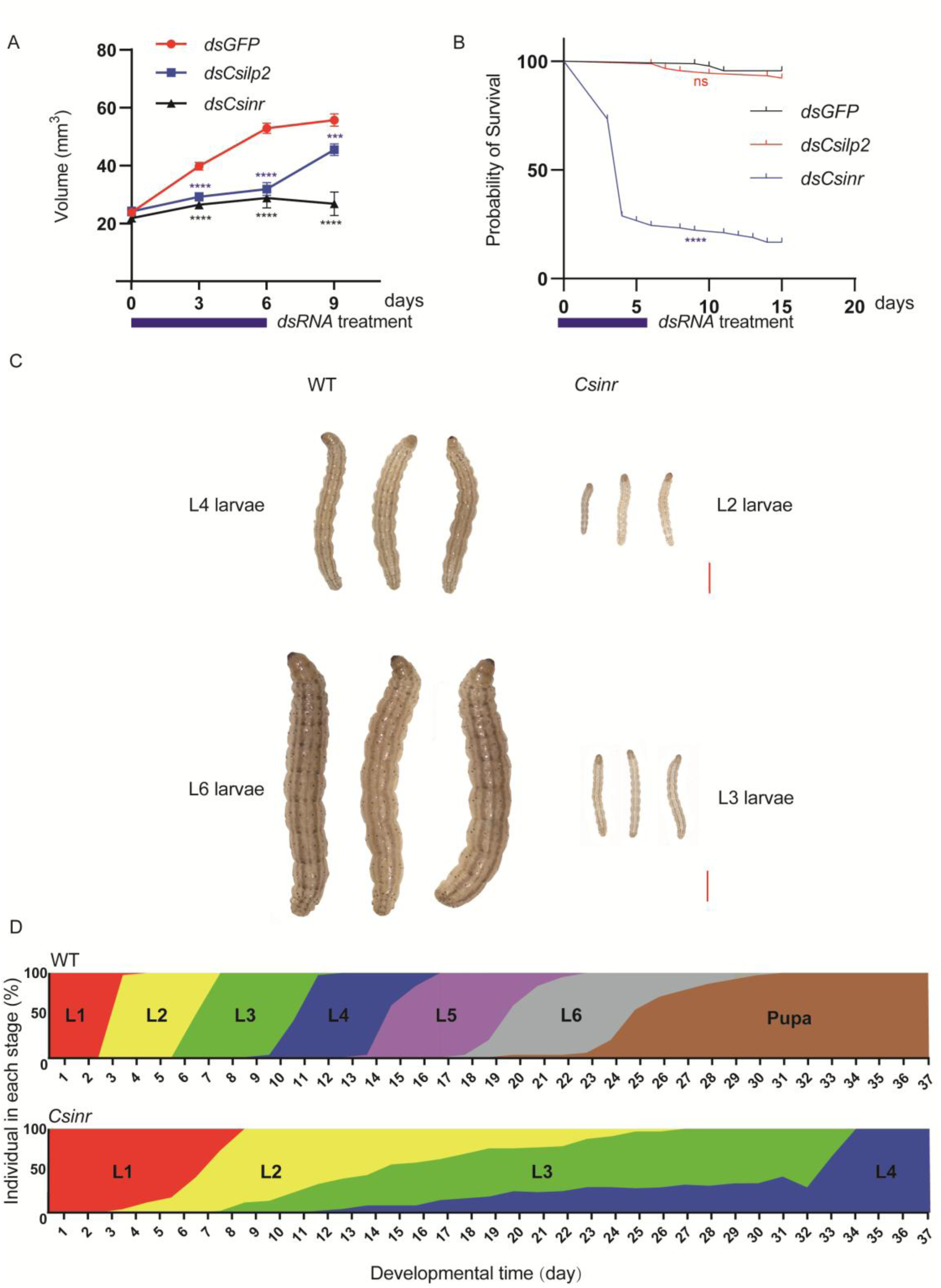
*Csilp2* and the ILP receptor, *Csinr,* regulates development and body size. (A) Body volume decrease in *C. suppressalis* larvae after knockdown of *Csilp2* and *Csinr* by feeding with *Csilp2* or *Csinr* dsRNA at stage early 5^th^ instar larvae. Two-way ANOVA and Šídák’s multiple comparisons test were used for data analysis, (*, *p* <0.05; ***, *p* < 0.001; ****, *p* < 0.0001; ns, no significant). (B) Survival rate of *Chilo suppressalis* larvae after feeding with *Csilp2* or *Csinr* dsRNA at stage early 5^th^ instar larvae. Statistics were performed using the Log-rank (Mantel-Cox) test; (*, *p* <0.05; ***, *p* < 0.001; ****, *p* < 0.0001; ns, no significant). (C) *Csinr* mutant larval phenotypes. Scale bar: 2 mm. (D) The stages of larval development in WT and mutants (*Csinr*) (L1, first instar; L2, second instar; L3, third instar; L4, fourth instar; L5, fifth instar; L6, sixth instar; Pupa).

**Fig. S11.**
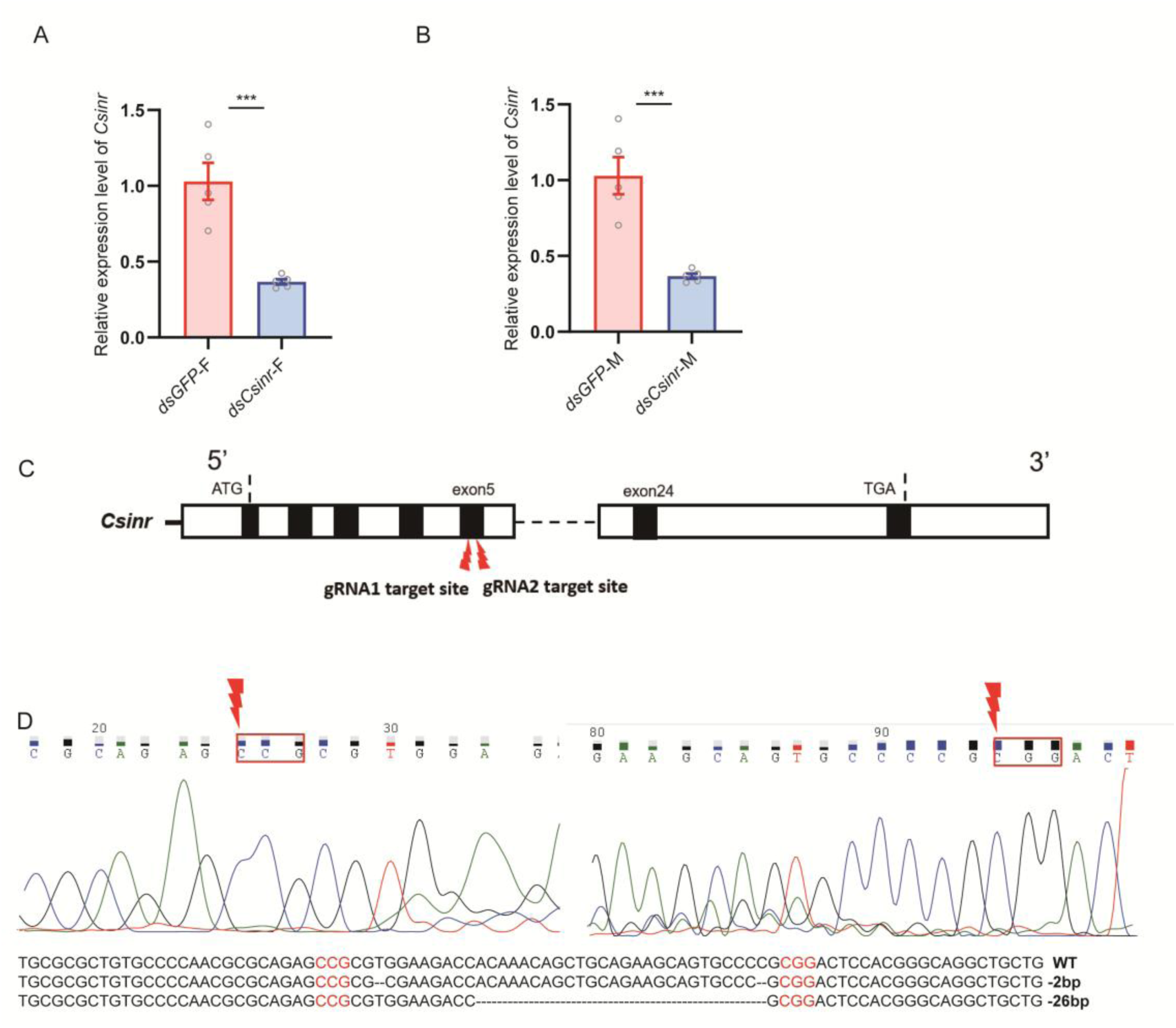
Validation of *Csinr* RNAi mediated knockdown and CRISPR/Cas9-mediated knockout of the *Csinr* gene in *Chilo suppressalis*. (A and B) Validation of *Csinr* RNAi silencing efficiency in both females and males by q-PCR. (C) Schematic diagram of gene sequences and two sgRNA target sites. Representative chromatogram of direct sequencing of PCR products of G0 individuals indicating that the two sgRNA sites were successfully cleaved. Green boxes indicate the PAM sequence. The position of the red lightning indicates the Cas9 protein cleavage site. (D) PAM sequences are highlighted in red. *Csinr* mutant sequences were confirmed by cloning and sequencing. Dashed lines represent the deleted bases, and inserted bases are displayed in lower case. The net change in length is marked at the right of each sequence (−, deletion; +, insertion). Statistical annotation: Data are presented as mean ± SEM. Student’s t-test. (*, *p* <0.05; ***, *p* < 0.001; ****, *p* < 0.0001; ns, no significant).

### Female-specific Csdsx isoforms directly bind and transcriptionally activate the Csilp2 promoter

We next asked whether *Csdsx* regulates *Csilp* expression, since our data indicated that the *Csdsx^E3^* mutation reduces larval size (volume) in *C. suppressalis*. We collected fourth- to sixth-instar *Csdsx^E3^* mutant larvae, with same-stage wild-type (WT) larvae serving as controls. Using qPCR, we demonstrated that WT females exhibited significantly higher *Csilp2* expression than males during both fifth and sixth instars, and interestingly, the *Csilp2* expression in *Csdsx^E3^* mutant females decreased significantly to levels comparable with males (Fig. 4A). Using *dsx*-RNAi during the fifth-instar larval stage, generated a similar result (Fig. 4B). Analysis of the promoter of *Csilp2* revealed eight potential *doublesex* binding cis-regulatory elements (CREs) located at -410 bp, -464 bp, -614 bp, -1300 bp, -1342 bp, -1644 bp, -1837 bp, and -2174 bp within the 2230 bp upstream region (Fig. 4C). The cloned *Csilp2* promoter fragment was subcloned into the pGL3-basic luciferase reporter vector. Concurrently, the full length coding sequences of the three *Csdsx* transcript variants were individually cloned into the pIB/V5-His expression vector (Fig. 4C). The recombinant plasmids were co<u>-</u>transfected into Sf9 (*Spodoptera frugiperda*) cells, with co-transfection of the empty vector serving as a negative control (Fig. 4C). Luciferase assays showed that transfection with the female<u>-</u>specific *dsx* splice variants significantly increased luciferase activity compared to the control, whereas transfection with the male<u>-</u>specific *dsx* variant significantly decreased luciferase activity (Fig. 4D). Consistent with the Luciferase assays results, electrophoretic mobility shift assays (EMSAs) were performed and the results showed that the DsxF1 protein strongly bound to CREs at -464 bp, -614 bp, -1300 bp, -1342 bp, -1644 bp, -1837 bp, and -2174 bp (Fig. 4E and F). DsxF2 protein strongly bound to CREs at -464 bp, -614 bp, -1300 bp, -1342 bp, -1644 bp, and 1837 bp (Fig. 4G and H). It appears that DsxM does not directly inhibit Csilp2 expression because of an obvious difference in its DNA-binding affinity compared to DsxF (Fig. 4I and J). Thus, based on these results, *Csdsx* appears to exert a sex-specific regulatory effect on *Csilp2* expression. These experiments therefore support that *Csdsx* regulates sex-specific growth via insulin signaling and does so by direct upregulation of *Csilp2* in female insects.

**Fig. 4.**
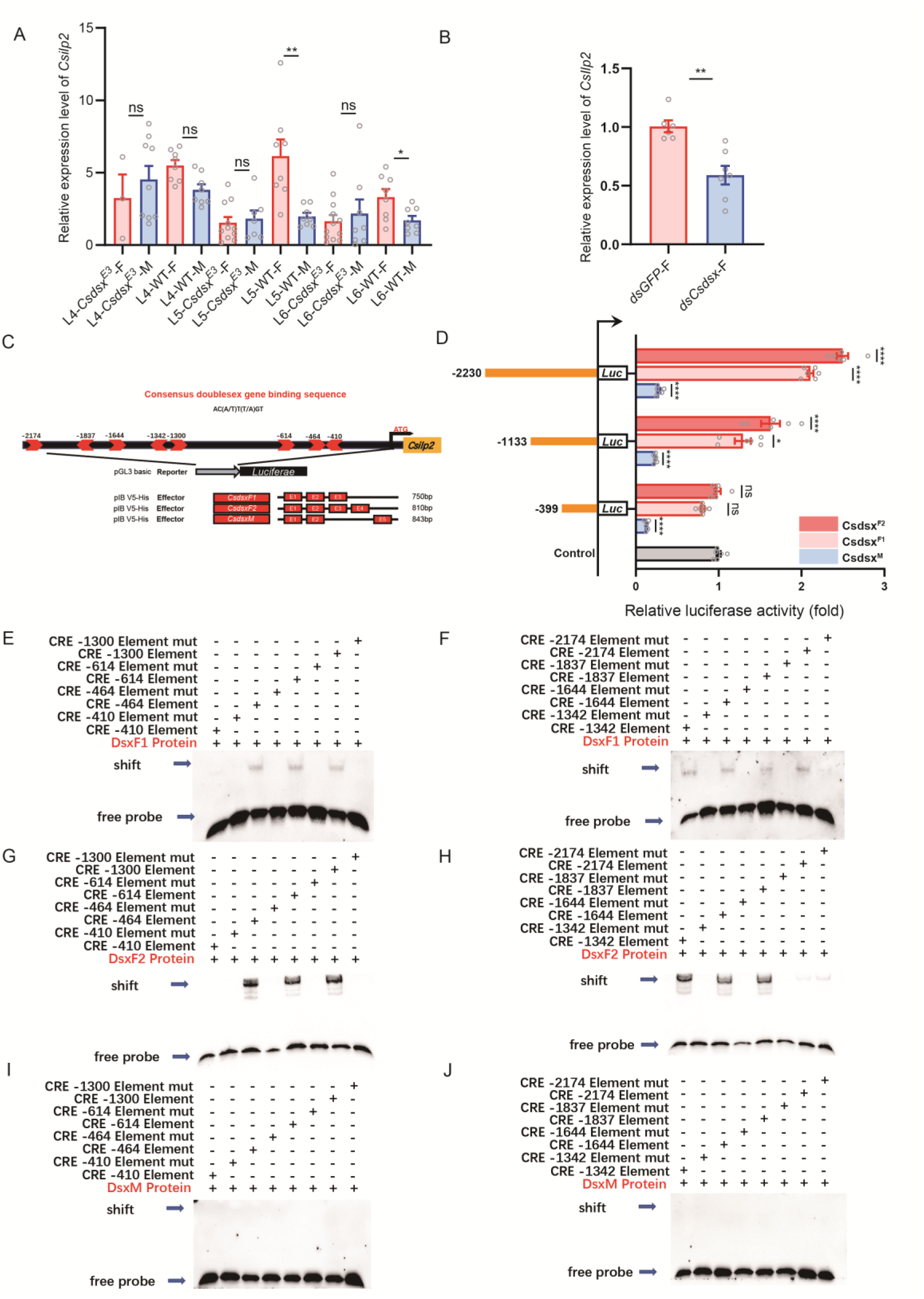
***Csdsx* regulates *Csilp2* expression.** (A) Expression analysis of *Csilp2* in female larvae of *Csdsx^E3^* mutants across different instar stages shows *Csilp2* expression in *Csdsx^E3^* mutant females decreased. Experimental samples: L4*-dsxE3-*F: Female *dsx^E3^* mutant 4th instar larvae; L4*-dsx^E3^-*M: Male *dsx^E3^* mutant 4th instar larvae; L4-WT-F: Female wild-type (WT) 4th instar larvae; L4-WT-M: Male wild-type (WT) 4th instar larvae. (B) Knockdown of *Csdsx* results in downregulation of *Csilp2* gene in females. (C) Schematic diagram of the *Csilp2* promoter constructs targeting different *Csdsx* splice variant vectors. (D) Differences in luciferase activity between the *Csilp2* promoter and different *Csdsx* splice variants. EMSA analysis of DsxF1 (E, F), DsxF2 (G, H) and DsxM (I, J) binding to the DNA fragments of *Csilp2* promoter. DNA binding activities of Dsx protein on each eight biotin-labeled doublesex CREs. “mut” indicates the mutant motif within the doublesex CREs. “+” indicated the presence of the sample in the lane; “-” indicated its absence. The black arrows point to the shifted bands and free probe. Statistical annotation: Data are presented as mean ± SEM. Student’s t-test. (*, *p* <0.05; ***, *p* < 0.001; ****, *p* < 0.0001; ns, no significant).

**Fig. 5.**
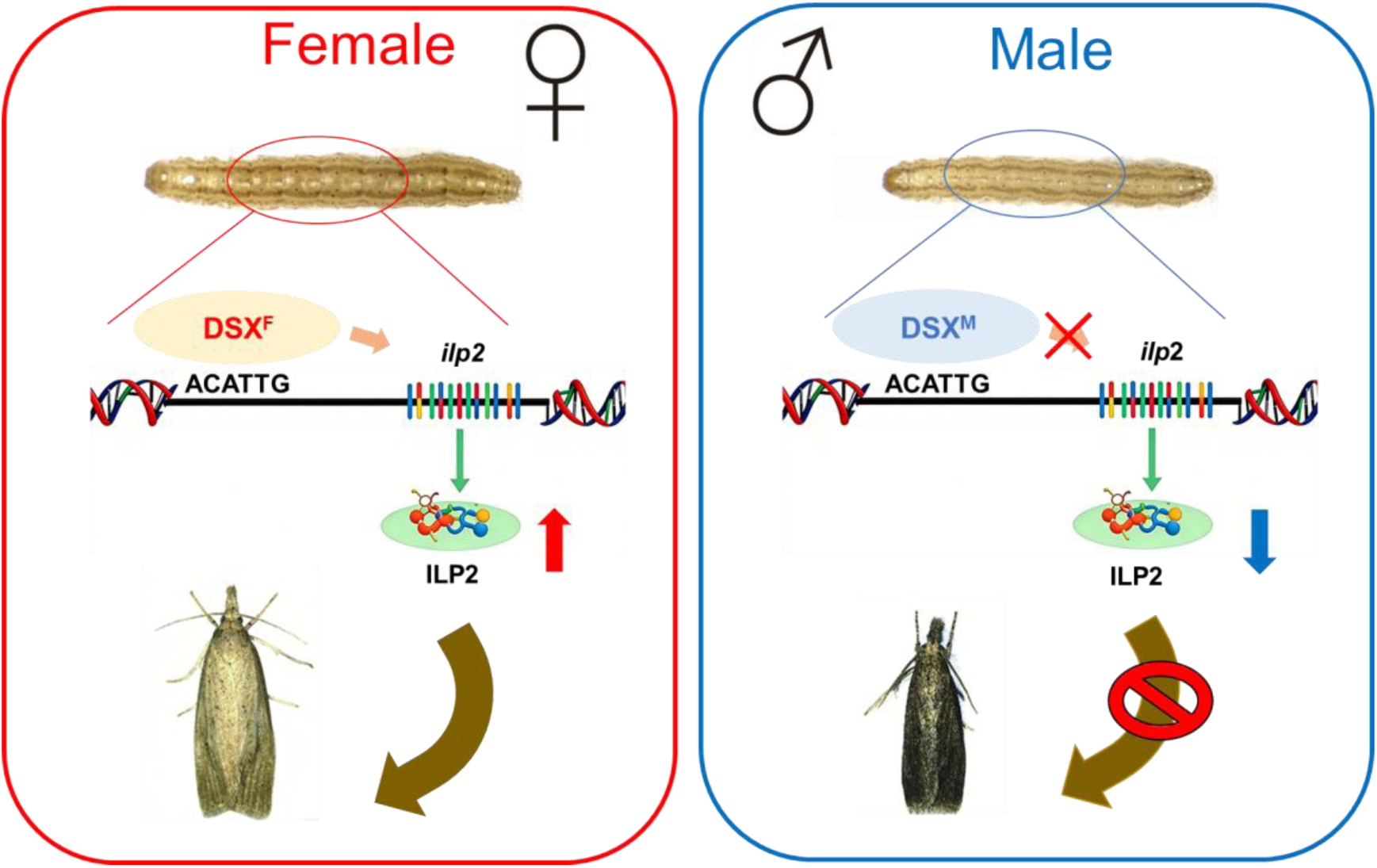
Proposed model for the underlying mechanisms of SSD in *C. suppressalis*. In *C. suppressalis*, the SSD is initiated in late fifth-instar larvae. The female-specific *Csdsx* isoform drives sex-biased body growth via transcriptional activation of the insulin gene *Csilp2* through its promoter binding sites. The resulting increase in ILP2 signaling is required for initiation of the SSD.

## Discussion

Our study shows that the sexual size dimorphism (SSD) in *C. suppressalis* does not arise in the egg (embryo) or early larval stages (L1–L4), but gradually emerges after L5 and is established at late L5. In female larvae the late-instar development is prolonged and displays a higher growth rate than males. Larval growth peaks at late L5, with male-to-female volume ratio reaching 1.3 by late L6. The absence of sex difference in the head capsule width suggests that the SSD is restricted to the thorax and abdomen, indicating a segment-specific growth regulation. The sex-determination gene *doublesex* (*dsx*), which is a crucial component of the insect sex determination pathway and plays a key role in insect sex differentiation and development (<u>Saccone, 2022</u>), is key to this SSD in *C. suppressalis*. We observed that knockout of the female-specific exon 3 of the *Csdsx* gene reduces pupal size and eliminates pupal SSD, while downregulating *Csdsx* in early L5 larvae eliminates SSD in early L6 and pupae. Furthermore, RNA interference-mediated reduction of *Csdsx* expression in early 5th instar larvae also eliminated the SSD in the early 6th instar and pupal stages.

Notably, transcriptomic analysis revealed that during the critical 5th and 6th instar stages, the gene encoding insulin-like peptide 2 (*Csilp2)* is highly expressed in late-instar female larvae, compared to males. Knockout of *C. suppressalis Csilp2* using CRISPR/Cas9 technology resulted in a prolonged duration of larval development and eliminated the SSD at both the larval and pupal stages. Importantly, the expression level of *Csilp2* was significantly decreased in female *Csdsx* E3 mutants. Finally, we identified a *dsx* binding site in the promoter region of *Csilp2*, and found that the female-specific *Csdsx* transcript exerts a promoting effect on the expression of *Csilp2* by targeting this promoter region. Thus, we identified a link between *dsx* and signaling with a specific ILP that regulates sex-specific growth of the female body during late larval development. Whereas ILPs have been shown earlier to regulate body growth in insects (<u>Brogiolo et al., 2001</u>; <u>Millington and Rideout, 2018</u>; <u>Okamoto and Mizoguchi, 2025</u>; <u>Texada et al., 2020</u>), this is the first demonstration of a sex-determination gene in regulation of a specific ILP to mediate sex-specific body growth.

Previous studies have indicated roles of *Dsx* in regulating sex-dimorphisms in the formation of body color, pigmentation patterns, and other sexual traits through distinct molecular mechanisms (<u>Guo et al., 2025b</u>) and here we add an effect on the SSD in *C. suppressalis*. We show that female-specific *dsx* splice variants are highly expressed in *C. suppressalis* during the period of rapid body growth in larvae, and we reveal an increased tissue-specific expression in head, fat body, and Malpighian tubules. Especially interesting is the *Csdsx* expression in the fat body, since this is also the major site where ILP2 is produced. Thus, *Csdsx* is likely to regulate the female-specific body growth by increasing ILP2 production and release from the fat body during the late larval stages and acting on progenitors of adult tissues.

Organismal growth is regulated by insulin/IGF signaling (IIS) in insects such as *Drosophila melanogaster* and *Bombyx mori* (reviewed in (<u>Brogiolo et al., 2001</u>; <u>Koyama and Mirth, 2018</u>; <u>Nässel and Broeck, 2016</u>; <u>Okamoto and Mizoguchi, 2025</u>; <u>Tennessen and Thummel, 2011</u>; <u>Texada et al., 2020</u>). In *Drosophila* insulin/IGF signaling (IIS) regulates organismal growth during both feeding (larval) and non-feeding (pupal) stages, although the larval growth is the most prominent (<u>Brogiolo et al., 2001</u>; <u>Ikeya et al., 2002</u>; <u>Okamoto and Mizoguchi, 2025</u>; <u>Okamoto et al., 2009b</u>; <u>Slaidina et al., 2009</u>; <u>Texada et al., 2020</u>). During the larval feeding stages *Drosophila* ILPs (DILPs), especially DILP2, produced by a set of median neurosecretory cells designated IPCs (insulin producing cells) are known to regulate growth (<u>Grönke et al., 2010</u>; <u>Ikeya et al., 2002</u>; <u>Okamoto and Mizoguchi, 2025</u>). On the other hand, during the non-feeding pupal stage, fat-body derived DILP6 is the main mediator of growth in *Drosophila* (<u>Okamoto et al., 2009b</u>; <u>Slaidina et al., 2009</u>) and also in *Bombyx* a fat body derived ILP regulates growth in the pupal stage (<u>Okamoto et al., 2009a</u>). To a lesser extent also the IPC-derived DILP1 regulates growth during non-feeding stages in *Drosophila* (<u>Grönke et al., 2010</u>; <u>Liao et al., 2020</u>; <u>Liu et al., 2016</u>). The fat body-derived ILPs in *Drosophila* and *Bombyx* are both related to insulin-like growth factors (IGF) of mammals (<u>Okamoto et al., 2009a</u>; <u>Okamoto et al., 2009b</u>; <u>Slaidina et al., 2009</u>). In *C. suppressalis*, however, we found that the fat body-derived ILP2 (CsILP2), which regulates sex-specific growth during feeding in the late larval development, is related to *Drosophila* DILP2 and not DILP6 or IGF. This is interesting to note while IIS dependent growth during feeding stages is directly reliant on nutrient intake, whereas growth during the pupal stage relies on nutrients stored during larval stages (<u>Liu et al., 2016</u>; <u>Okamoto and Mizoguchi, 2025</u>; <u>Okamoto et al., 2009b</u>; <u>Slaidina et al., 2009</u>; <u>Texada et al., 2020</u>). Thus, the action of the two fat-body derived ILPs differ between *Drosophila* and *C. suppressalis*, CsILP2 acts during feeding in the late larval stage and DILP6 during pupal development.

Importantly, growth regulation in insects relies both on ILPs and their interactions with signaling by ecdysone and juvenile hormone (<u>Koyama and Mirth, 2018</u>; <u>Okamoto and Mizoguchi, 2025</u>; <u>Texada et al., 2020</u>). The latter signals are critical since they regulate timing of developmental events and determine the duration of the larval feeding stage and thus affect growth (<u>Okamoto and Mizoguchi, 2025</u>; <u>Tennessen and Thummel, 2011</u>; <u>Texada et al., 2020</u>), and prolonged larval feeding leads to increased size. Indeed, we found that the late larval instars of females feed several days longer than the males do (Table S1), which likely contributes to the increased size. Thus there may be a differential involvement of ecdysone signaling to increase the feeding duration in the 5^th^ and 6^th^ instars in females. Mechanistically, ILP action on tissues involves the canonical target of rapamycin (TOR) pathway, which is nutrient dependent and regulates translational processes (via ribosomes) and growth, but ILPs also interact with FOXO in growth regulation (<u>Garofalo, 2002</u>; <u>Koyama and Mirth, 2018</u>; <u>Okamoto and Mizoguchi, 2025</u>; <u>Texada et al., 2020</u>). While the SSD detected in *C. suppressalis*, is mediated by increased activation of the InR and TOR pathway in growing tissues, ILP2 seems not to act on developing head structures where no SSD is seen, or exerts only negligible effects.

In *C. suppressalis*, both the *dsx* gene and the insulin-signaling pathway (specifically ILP2) are involved in regulating the SSD. To investigate how *dsx* regulates insulin signaling in *C. suppressalis*, a dual-luciferase reporter assay system was deployed, this system enables direct visualization of the regulatory effect of specific transcription factors on the promoter activity of target genes. We found that female-specific splice variants of *Csdsx* activate the *Csilp2* promoter. This indicates that *Csdsx* regulates the transcriptional activity of *Csilp2* by directly binding to the *dsx*-specific cis-regulatory elements within the *Csilp2* promoter.

The role of Dsx in *C. suppressalis* in regulating the whole body SSD shown here is novel. Other sex-dimorphisms in growth of specific organs have been demonstrated previously. In dung beetles (*Onthophagus*), the male-specific *dsx* splice variant (*dsxM*) drives the nutrition-dependent growth of male horns, whereas the female-specific splice variant (*dsxF*) inhibits horn formation in females (<u>Kijimoto et al., 2012</u>). It was also shown that in *C. suppressalis Csdsx* is involved in regulating antenna width, adult wing color, adult external genitalia, adult internal reproductive system, and fecundity (<u>Guo et al., 2025a</u>). Collectively, these findings indicate that *dsx* achieves specific and precise regulation of sexual dimorphism in growth of various insects by sex-specific splicing of the *dsx* gene. There are also reports on other sex-determination genes that are involved in establishing a SSD in insects. In *Drosophila*, *sexless* (*Sxl*)-expressing neurons in the brain play a crucial role in regulating early size differences between males and females (<u>Sawala and Gould, 2017</u>). Additionally, expression of the *transformer* (*tra*) gene in the female fat body of *Drosophila* promotes growth by signaling to the IPCs in the brain to induce ILP secretion (<u>Rideout et al., 2016</u>). Finally, in *Onthophagus taurus*, studies on horn development have demonstrated that *Dsx* participates in the nutrition-sensitive regulation of horn growth, ultimately leading to sexual dimorphism in horn morphology (<u>Kijimoto et al., 2009</u>).

In conclusion, we found that in *C. suppressalis* (1) the SSD originates during the late 5^th^ instar larval stage, (2) a female-specific splice form of the sex-determination gene *Csdsx* plays a critical role in the sex-specific body growth, (3) the SSD depends on signaling with fat body derived ILP2 (encoded by *Csilp2*) and (4) *Csdsx* regulates production of CsILP2 by activating binding sites on the *Csilp2* promoter. Thus, we have identified a novel pathway that regulates sex-specific body size during development.

## Materials and Methods

### Insects

The strain of *C. suppressalis* was provided by Dr. HAN Lan-Zhi of the Chinese Academy of Agricultural Sciences in 2018 and reared in the laboratory. Larvae were fed on artificial diet (<u>Han et al., 2012</u>) Rearing conditions were 28 ± 1 °C, 70–80% relative humidity (RH) and a 16 h:8 h, light : dark photoperiod. Larvae were collected in plastic petri dishes for pupation. Moths were transferred to insect cages with a RH of 85%-90% and fed with 10% honey water.

### Gene cloning and sequence analysis

We used the NCBI database and BLAST programs for sequence alignment and analysis. Then we used EditSeq to predict Open Reading Frames (orfs). The primers were designed by tools in NCBI. The primer sequence information is described in (Table S2). According to the manufacturer’s instructions, total RNA was extracted from pupae using Trizol reagent (Invitrogen, Carlsbad, CA, USA). RNA quality was assessed by 1.2% agarose gel electrophoresis and the concentration of RNA was measured using a spectrophotometer (Nanodrop 2000, thermo, Shanghai, China). The complementary DNA was synthesized by HiFiScript cDNA Synthesis Kit (Proteinssci, Shanghai, China) following the manufacturer’s protocol. Each 25 μL PCR reaction included 12.5 μL 2 × phanta Max Master Mix (Vazyme, Nanjing, China), 1 μL of forward primer (10 μmol/L), 1 μL of reverse primer (10 μmol/L), 2 μL of cDNA, and 8.5 μL ddH2O. The specific reaction procedure was as follows: 95 ◦C for 3 min, followed by 35 cycles of 95 ◦C for 15 s, 57 ◦C 15 s, 72 ◦C for 1 min, and final extension at 72 ◦C for 10 min. The PCR amplification product was purified with Gel Extraction Kit (Omega, USA) and cloned into pClone007 Blunt Simple vector (TSINGKE, Beijing, China) for sequencing.

Multiple alignments of full-length sequences were performed with the ClustalW Multiple Alignment program in BioEdit. Phylogenetic relationships were inferred using the maximum likelihood method with the Whelan and Goldman model (<u>Whelan and Goldman, 2001</u>). Phylogenetic tree was constructed using MEGA 12 software with the Maximum Likelihood Method and bootstrapped with 1000 replications, Gamma parameters: 4 discrete categories (<u>Kumar et al., 2024</u>). Initial trees for the heuristic search were obtained automatically by applying neighbor-joining and BioNJ algorithms to a matrix of pairwise distances estimated using the JTT model, and then selecting the topology with a superior log likelihood value. Rates among sites choose Gamma Distributed With Invarant Sites (G+l). A discrete gamma distribution was used to model evolutionary rate differences among sites at 5 categories. The analysis of amino acid sequences with positions exhibiting less than 95 % site coverage eliminated (i.e., fewer than 5 % alignment gaps, missing data, and ambiguous bases were allowed at any position).

### Molecular sex identification

Molecular sex identification was measured as previously described(<u>Guo et al., 2025a</u>).

### Volume and weight of larvae and pupae

All images were captured using a ZEISS Stemi 2000-C stereomicroscope equipped with an integrated digital camera. Developmental time from hatching to pupation was recorded daily, beginning with the egg stage. Following egg hatching, individuals were photographed each day until pupation. Artificial diet was refreshed daily, and the timing of each molt was noted. On the second day after pupation, pupae were weighed and sexed. Larval body length and width, pupal length and width were measured from digital images using ImageJ software. Adult body length, body width, wing length, and wing width were measured on the second day post-eclosion. Image processing and figure preparation were performed using ImageJ and Adobe Illustrator. Larval and pupal volumes were calculated according to the formula for Drosophila melanogaster volume measurement: V=1/6π(L)(W)^2^(<u>Rideout et al., 2016</u>).

### Quantitative real-time PCR

The expression patterns in different developmental stages were studied using qRT-PCR. Third-instar larvae (n = 10-15), fourth-instar larvae (n = 5-10), fifth-instar larvae (n = 5-10), sixth-instar larvae (n = 5-10), wandering larvae (n = 5-10). Different development tissues were dissected from adult females and males (day-2 adults) for each sex, including head (n = 10), thorax (n = 5), legs (n = 10), abdominal epidermis (n = 5), wings (n = 10), gut (n = 10), fat body (n = 10), Malpighian tubules (n = 10), ovary (n = 5), testis (n = 10), reproductive system (n = 10), antenna (n = 20), copulatory pouch (n = 10), female accessory gland (n = 10). RNA extraction and cDNA synthesis were the same as for RT-PCR assay. The qRT-PCR analysis was performed with AceQ Universal SYBR qPCR Master Mix (Vazyme, Nanjing, China) using a QuantStudio 5 Real Time PCR System (Applied Biosystems, USA). The 20 μL reaction included 2 μL of 10-fold diluted cDNA. Four biological replicates were performed for each sample. Relative mRNA levels were calculated using the 2^−△△Ct^ method (<u>Livak and Schmittgen, 2001</u>). *Actin* and *G3PDH* were used as internal controls. Primers used in RT-PCR and qRT-PCR are shown in (Table S2).

### CRISPR/Cas9 genome editing

Based on the target sequence criteria of 5’-GGN18NGG-3’, the single guide RNA (sgRNA) was designed against target gene via the CRISPR gRNA Design tool-ZiFiT 4.2 (http://zifit.partners.org/ZiFiT/Disclaimer.aspx). The sgRNAs were synthesized *in vitro* using the GeneArt™ Precision gRNA Synthesis Kit (Thermo Fisher Scientific, Vilnius, Lithuania) according to the instructions of manufacturer. The Cas9 protein (TrueCut™ Cas9 Protein v2, Cat. NO. A36497) was purchased from Thermo Fisher Scientific (Shanghai, China).

Newly laid eggs were neatly lined on the glass slide with double-sided adhesive tape (approximately 1 h-old). The microinjection was performed using a microinjection system (InjectMan NI 2 microinjection system, Eppendorf, Hamburg, Germany). After Cas9 protein and gRNA were incubated for 30 min at 25 ℃, the mixture of Cas9 protein (300 ng/μL) and sgRNA (150 ng/μL) was co-injected into embryos. Injected embryos were incubated at 26 ± 1 ℃ and 70% ± 5% RH for hatching.

### Generation and characterization of mutants

To determine whether CRISPR/Cas9 successfully knocked out target gene, the genomic DNA fragments of target gene flanking the target regions. Genomic DNA was isolated from ten newly hatched larvae randomly(<u>Sun et al., 2023</u>). The PCR amplification products were run on a 1.2% agarose gel to verify that the mutation occurred. The PCR products were cloned into the pClone007 Simple vector (TSINGKE, Beijing, China) and sequenced using the specific primers (Table S2). Sequencing of the G0 mutants showed multiple peaks near the target site, suggesting mutations.

### RNA interference

The dsRNA template was generated as follows. The gene served as the PCR template. Gene-specific primers (Table S2), each bearing a T7 promoter sequence at the 5′ end, were designed to amplify the region common to both male and female *<u>dsx</u>* transcripts. PCR amplification was performed under standard conditions, and the resulting amplicons were purified using gel-extraction kit (E.Z.N.A.® Gel Extraction Kit) to yield clean cDNA templates.

In vitro transcription was carried out with the MEGAscript T7 Kit (Thermo Fisher Scientific) according to the manufacturer’s instructions. Each 20 µL reaction contained 200 ng of purified cDNA template, NTPs, T7 enzyme mix, and buffer, and was incubated at 37 °C for 16 h. After transcription, 1 µL of DNase I was added to degrade the DNA template; the mixture was gently mixed and incubated at 37 °C for 15 min, followed by enzyme inactivation at 65 °C for 10 min. To prepare samples for quality control, 0.5 µL of the transcription reaction was mixed with 9.5 µL of RNase-free water. Eight microliters of this dilution were evaluated by agarose gel electrophoresis to confirm the size and integrity of the dsRNA, and the remaining 2 µL were used to determine dsRNA concentration via UV spectrophotometry (NanoDrop 2000, Thermo Fisher Scientific). Final dsRNA aliquots were stored at – 80 °C until further use.

Microwave-assisted synthesis of CQD–dsRNA complexes was performed as follows. First, 9 mL of polyethylene glycol was mixed with 3 mL of deionized water. Separately, 100 mg of polyethyleneimine was dissolved in 2 mL of deionized water, and the combined solution was subjected to microwave irradiation at 800 W for 3 min to yield functionalized carbon quantum dots (CQDs). Pre-synthesized dsRNA was then suspended together with the functionalized CQDs in a sodium sulfate solution and incubated overnight at 4 °C to form CQD–dsRNA complexes (<u>Wang et al., 2020</u>).

For RNAi feeding assays, artificial diet was cut into rectangular pellets measuring 5 mm × 5 mm × 3 mm (0.2 g each). Approximately 5 µg of dsRNA–CQD suspension was used to coat each diet pellet, which was then fed to cohorts of ten newly molted fifth-instar larvae. Diet pellets were refreshed daily for six consecutive days (<u>Wang et al., 2020</u>). On day 6, twelve larvae were collected for total RNA extraction, sex determination, and assessment of gene-silencing efficiency.

### RNA-seq and data analysis

24 male and 24 female larvae were collected on day 3 of the fourth, fifth, and sixth instars. Following sex determination, five individuals per sex from each instar were pooled for total RNA extraction. Extracted RNA samples were immediately flash-frozen on dry ice and shipped to Novogene Co., Ltd. (Beijing) for downstream analysis. Five biological replicates were prepared. RNA extraction was performed using the TRIzol reagent (Invitrogen) following the manufacturer’s protocol. RNA quality and quantity were assessed using an Agilent 2100 BioAnalyzer (Agilent, USA), and RNA integrity was confirmed through electrophoresis on a 1% agarose gel. RNA-seq libraries construction, sequencing and assembly of transcriptome reads was performed by Novogene with Illumina HiSeq2000 platform (Novogene Bioinformatics Technology Co.Ltd, Beijing, China) and Trinity (v2.4.0), respectively.

The clean reads were generated by removing adapters, poly-N, and low-quality reads from the raw data using fastp algorithm (v0.12.4). Sequences were aligned to the *Chilo suppressalis* genome (GenBank: GCA_004000445.1) using Hisat2 v2.0.5. Differential expression analysis was performed using the DESeq2 R package (1.16.1). Log2fold change >1 and FDR < 0.01 were used as screening criteria to screen for differentially expressed genes. The cluster Profiler R package was used for Gene Ontology (GO) enrichment analysis and KEGG pathway analysis. FDR-adjusted multiple tests were added to the hypergeometric test.

### Vector construction and transfection

Transcription factor binding sites in the 5′ flanking region of the *Csilp2* start codon were predicted and analyzed using the JASPAR (http://jaspar.genereg.net) and ALGEN PROMO (http://alggen.lsi.upc.es/cgi-bin/promo_v3/promo/promoinit.cgi?dirDB=TF_8.3) online platforms.

Using the *Chilo suppressalis* genome, we designed primers for the *Csilp2* gene to facilitate cloning of this region. Primers were designed within 100 bp downstream of the start codon, and the forward and reverse primers contained approximately 1000 bp of sequence. GeneDoc software was used to compare the obtained sequences. Primers are reported in (Table S2). Single-head DNA was extracted from pupae of *Chilo suppressalis*. The PCR reaction mix was prepared with the following components: 1 μL of forward primer, 1 μL of reverse primer, 2 μL of DNA template, 12.5 μL of 2 × Rapid Taq Master Mix, and double-distilled water (ddH₂O) to a final volume of 25 μL. The PCR program included an initial denaturation at 95 ◦C for 3 min, followed by 35 cycles of denaturation at 95 ◦C for 15 s, annealing at 57 ◦C for 15 s, and extension at 72 ◦C for 3 min. The reaction concluded with a final extension at 72 C for 5 min and was held at 4 ◦C.

Cloning primers for the transcription factors were designed based on three *C. suppressalis dsx* splice variants (*dsxF1, dsxF2*, and *dsxM*). For each target, gene-specific primers were synthesized to amplify the full coding region, with overhangs corresponding to the KpnI and XbaI sites of the PIBV5-His expression vector. Specifically, the 5′ and 3′ ends of each primer incorporated homology arms matching the vector sequence flanking the KpnI and XbaI restriction sites. Using *Chilo suppressalis* cDNA as template, the desired fragments were PCR-amplified with a high-fidelity DNA polymerase. The PIBV5-His vector was then linearized by double digestion with KpnI and XbaI, and the purified PCR products were ligated into the vector. Recombinant plasmids were transformed into competent Escherichia coli cells, and positive clones were screened by colony PCR. Confirmed clones were cultured, and plasmid DNA was submitted for sequence verification.

Differences in the activity of *Csilp2* gene promoter sequences were analyzed using dual-luciferase reporter assays. The *Csilp2* promoter sequences were ligated into pGL3-Basic vector (Promega, USA). Sf9 cells were spread in cell culture plates (24 wells) at a density of 5 × 10^5^ cells in a volume of 1 ml/well. Then the cells were transfected with 1 μg promoter construct plasmid or expression plasmid and 0.1 μg pRL-CMV and 3 μl of Lipofectamine® 2000 Reagent transfection reagent in 200 μl of DMEM medium per well. Cells were lysed for fluorescence detection after 24 h, at least 3 replicate experiments were performed.

We also conducted an additional dual luciferase assay to test how variants in the promoter affected expressional activity. In this article, when describing promoter sequences, a negative sign (−) is used to indicate the position relative to the transcription start site. We designate the transcription start site as +1, with the negative sign representing a position upstream of the transcription start site (i.e., in the 5′ direction). For example, the −399 region is located 399 bases upstream of the +1 site.

We constructed different artificial promoter sequences by truncating the sequence from the 5′ end (further upstream from the gene) while retaining the sequence at the 3′ end (closer to the gene). The first truncation tested retained sequences from −2230 to −1 upstream, which we call the P (−2230/−1) sequences. The second truncation tested retained sequences from −1133 to −1 upstream, which we call the P (−1133/−1) sequences. The third truncation tested retained sequences from −399 to −1 upstream, which we call the P (−399/−1) sequences. As a control, we used the empty pGL3-Basic vector.

### Electrophoretic mobility shift assays (EMSA)

The gene *dsx* were cloned from the cDNA of *Chilo suppressalis.* to construct the plasmid. EcoR I and Hind III were used to ligate *dsx* gene fragments to the prokaryotic expression vector pET30 a to construct the recombinant plasmid pET30 a-dsx. After double enzyme digestion and identification with PCR, the recombinant plasmid was sent for sequencing. The sequence of the recombinant plasmid was verified. The recombinant plasmid was transformed into competent cells (*Escherichia coli* BL21, DE3), and expression of the protein expression was induced with IPTG. The suspension was sonicated, and protein expression was analyzed using 12% SDS-PAGE. The proteins were purified on a HIS Trap purification column.

Probes for dsx CRE -410, -464, -614, -1300, -1342, -1644, -1837, and -2174 were created (Table S2). These dsx CREs were predicted possible binding sites on the Csilp2 promoter according to the website https://jaspar.elixir.no/. These probes, with biotin-labeled (produced by EMSA Probe Biotin Labeling Kit, Beyotime, GS008, China), underwent annealing through a 10 minutes heating process at 95 °C, followed by gradual cooling to 25 °C. The purified proteins were then incubated with the biotin-labeled probes. The incubation process followed the guidelines provided in the EMSA/Gel-Shift Kit (Beyotime, GS009, China). To evaluate dsxF1, dsxF2, and dsxM ability to bind DNA, proteins were combined for 20 minutes. Subsequently, these mixtures were incubated to their respective probes for analysis. The resulting mixtures were separated using 6% (w/v) native polyacrylamide gels and underwent electrophoresis in TBE buffer (45 mM Tris-borate, 1 mM ethylenediaminetetraacetic acid, pH 8.3). Post-electrophoresis, the samples were transferred onto a nylon membrane (Biosharp BS-NY-45, China). The bands were then visualized using a chemiluminescence gel imaging system (Tanon Chemi Dog Ultra automatic luminescence imaging system).

## Statistics

We used the GraphPad Prism 10 software package to generate graphs and statistically analyze data. The Mann Whitney test was used to compare two columns. The Kruskal-Wallis one-way ANOVA test, followed by Dunn’s post test, was used to compare multiple columns of data. Student’s t test was used for pairwise comparisons. All data are presented as mean ± s.e.m. The sample sizes, number of replicates and statistical tests used for each experiment are stated in the figures or figure legends.

## ACKNOWLEDGMENTS

This work was supported by the National Key R & D Program of China (2025YFE0210000 to Shun-Fan Wu) and National Natural Science Foundation of China (Grant No. 32022011 & 32272576 to Shun-Fan Wu).

---

**Table S2.**
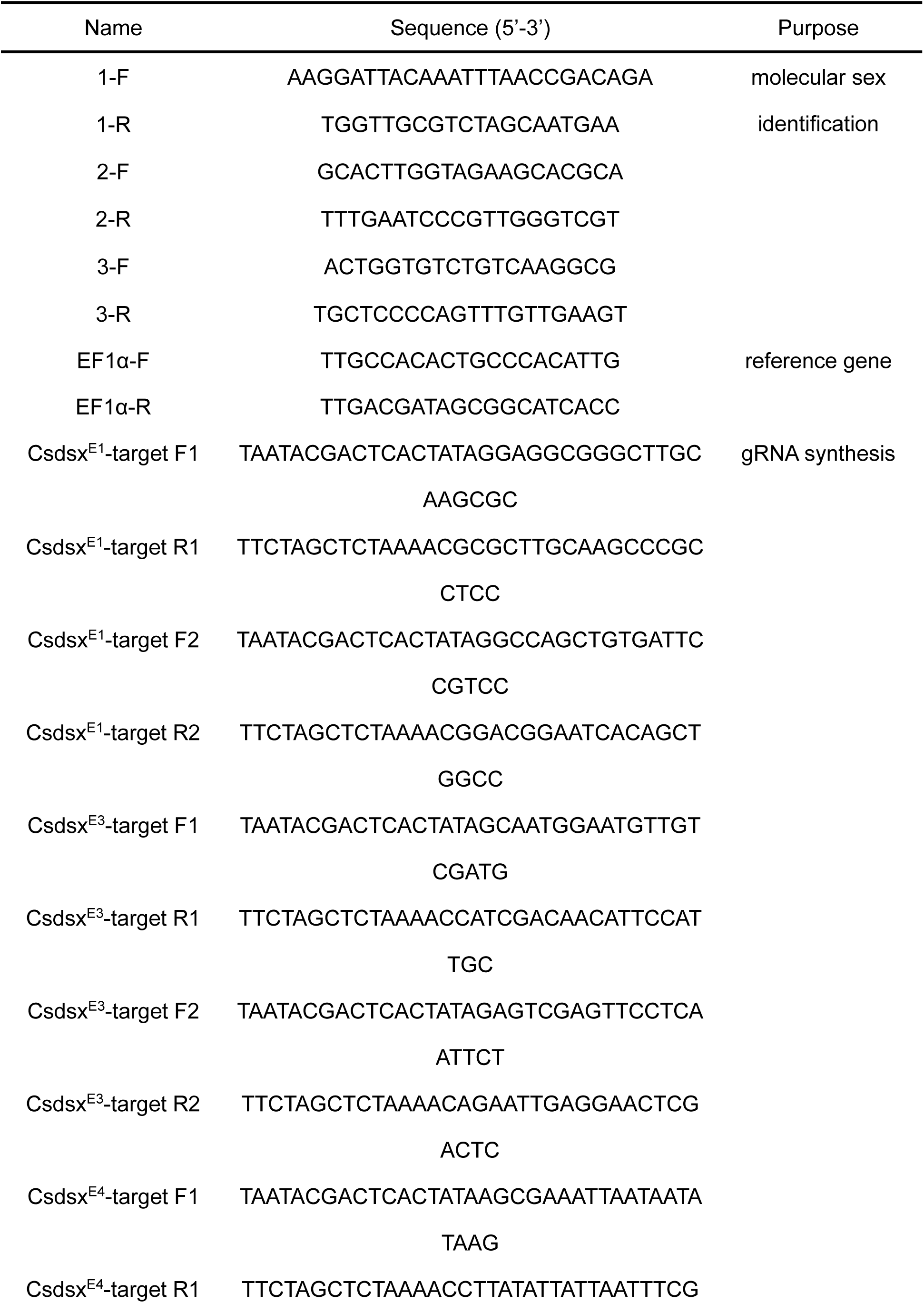
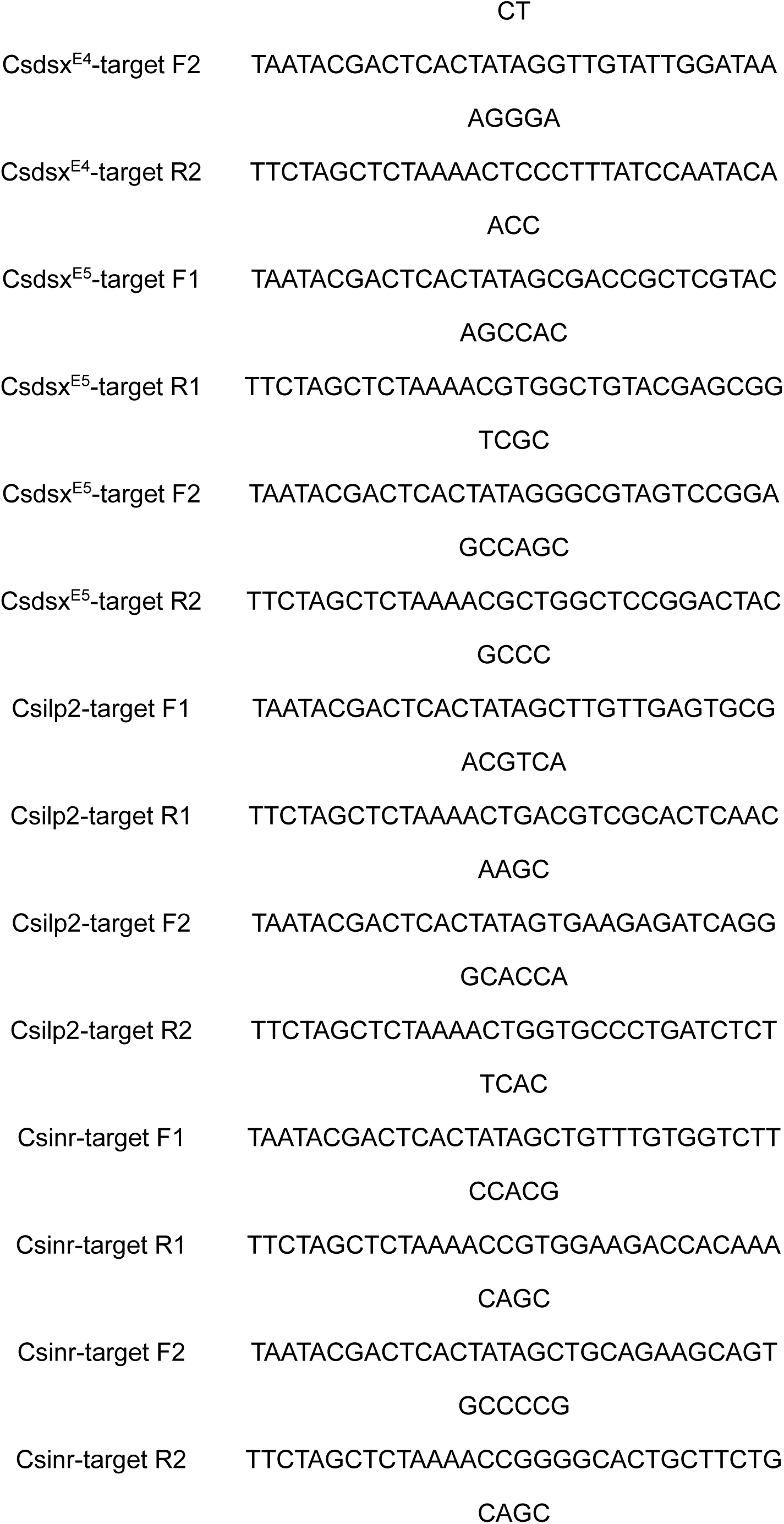
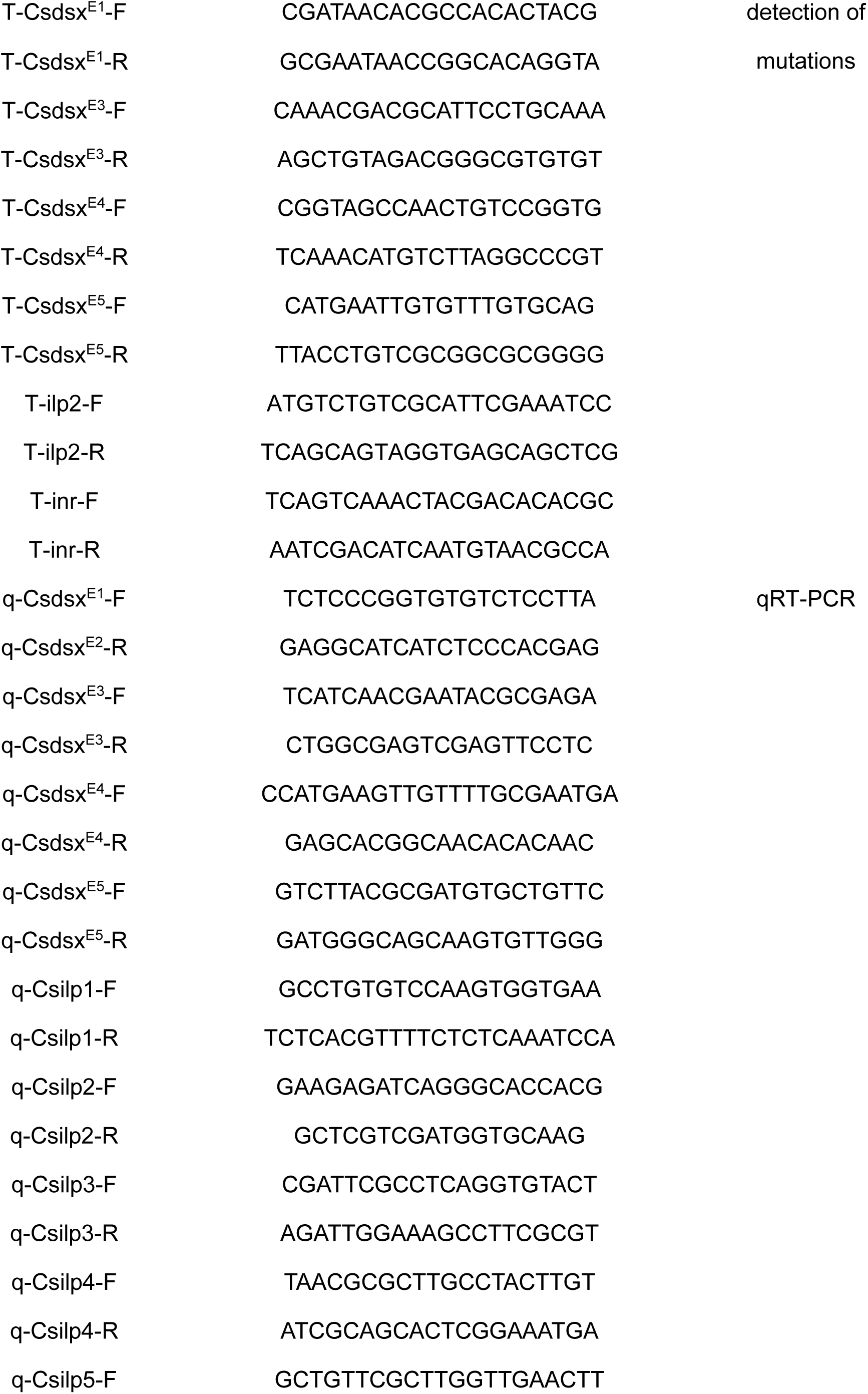
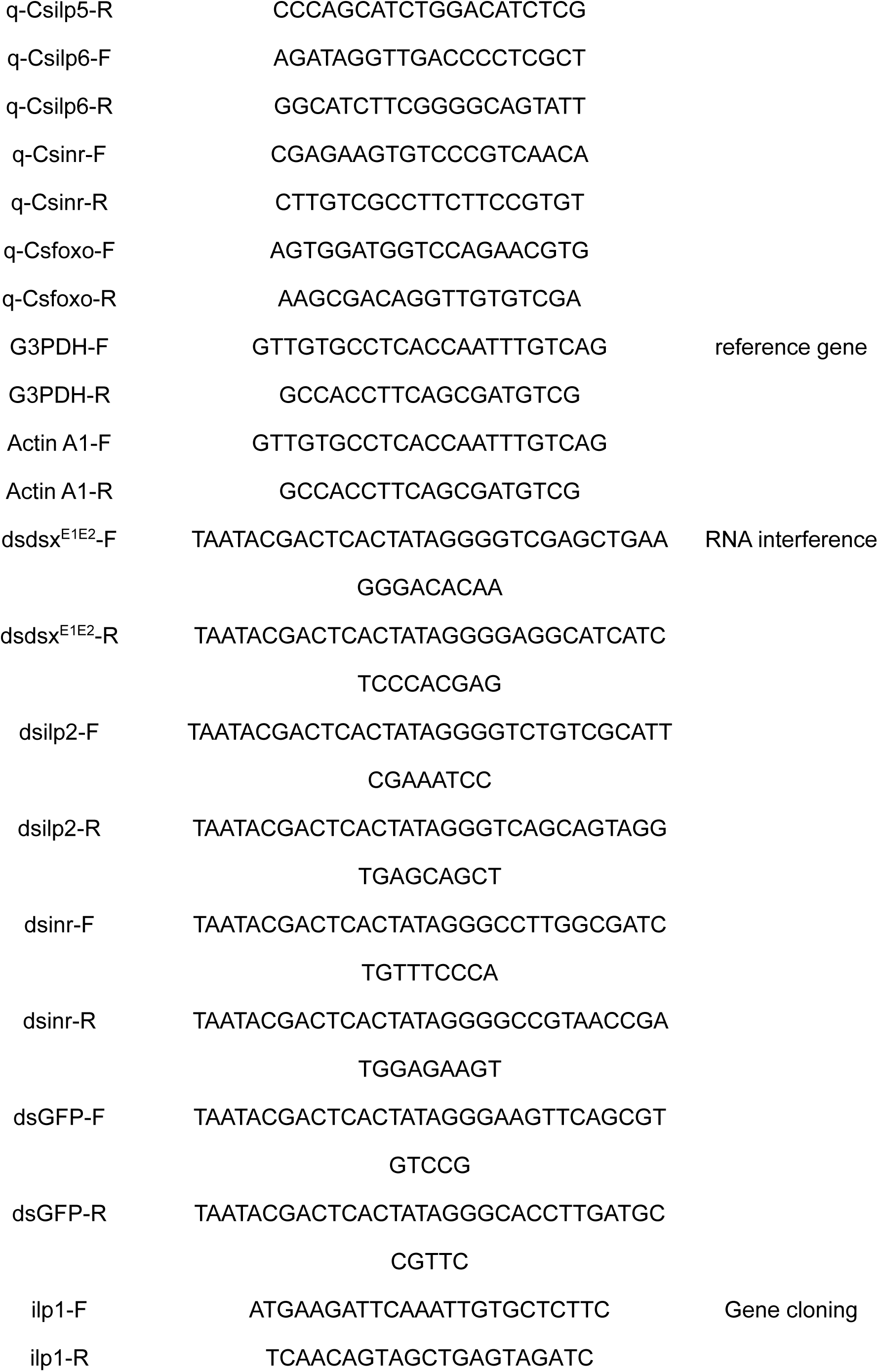
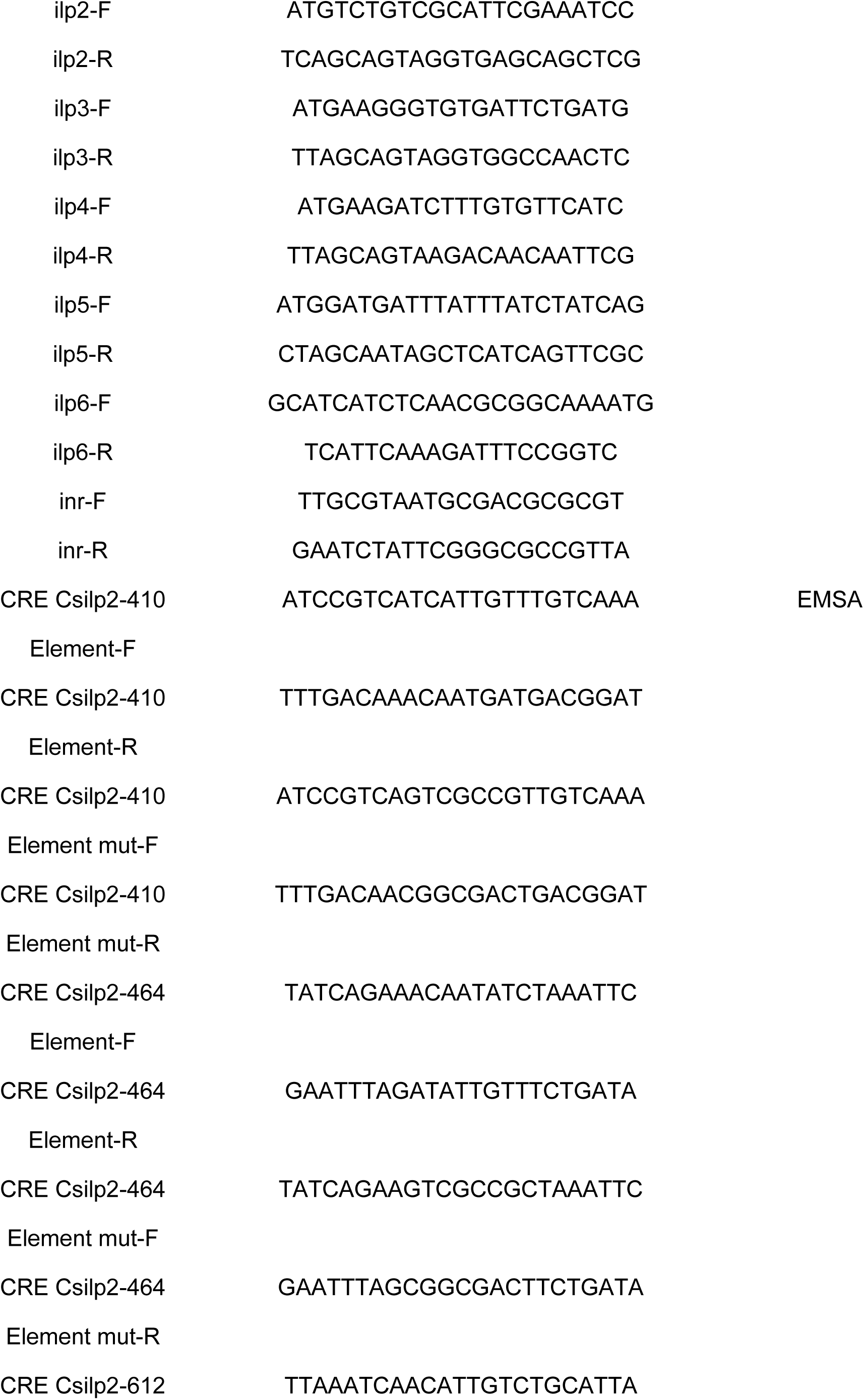
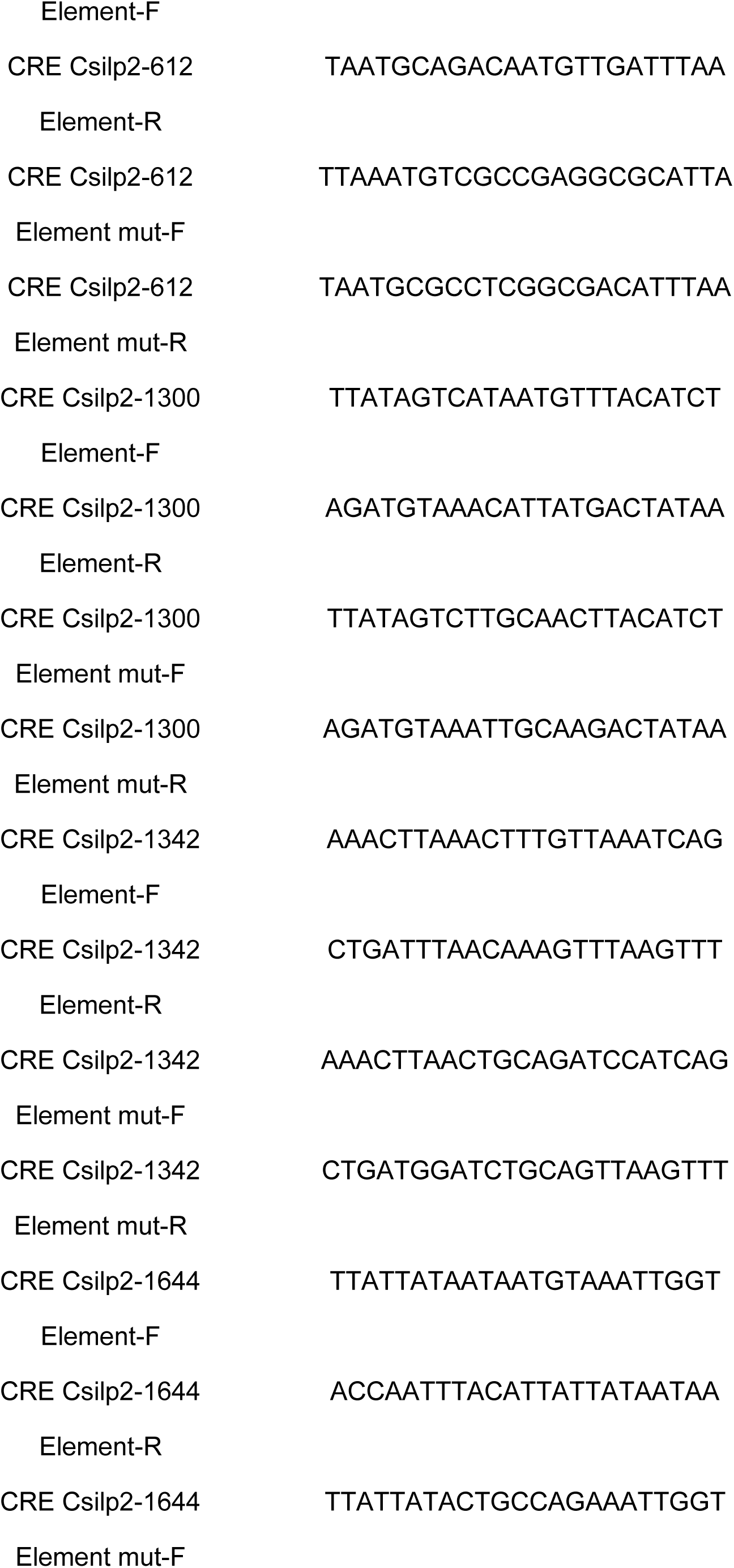
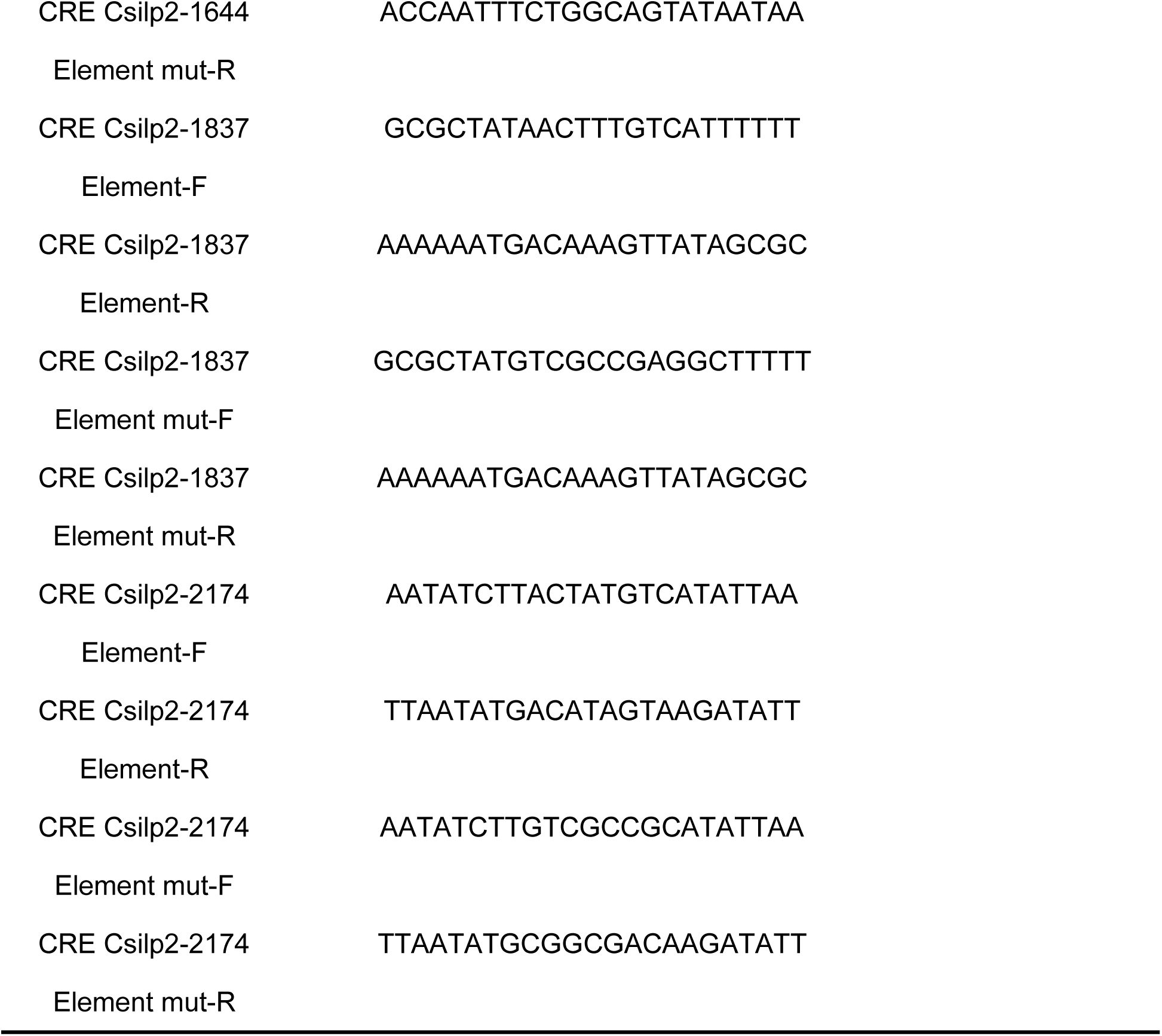
The primers used in this study.

